# Intrinsic antifungal activity of curli nanofibers expands the design space for programmable antimicrobial biomaterials

**DOI:** 10.64898/2026.08.18.745637

**Authors:** Nicolas Burns, Anna Kurowski, Hoda M. Hammad, Braden Ross, Matthew Bryant, Anna M. Duraj-Thatte

## Abstract

The rise of antifungal resistance and the limited number of available antifungal drug classes have created an urgent need for biomaterials capable of localized, programmable antifungal activity. Microbial extracellular protein nanofibers have emerged as versatile scaffolds for engineering functional biomaterials, yet whether these structural proteins possess intrinsic biological activities remains largely unexplored. Here, we engineered curli nanofibers displaying the antifungal peptide heliomicin and unexpectedly discovered that wild-type CsgA itself exhibits intrinsic antifungal activity against *Candida albicans*, reducing fungal viability by approximately 2 log units. Genetic fusion of heliomicin enhanced this intrinsic activity to a 3.5-log fungicidal reduction while preserving nanofiber self-assembly, hydrogel formation, mechanical properties, and 3D printability. Mechanistic analyses identified membrane disruption as the primary mode of action and linked the enhanced activity of Hel-CsgA to expansion of the cationic surface of CsgA. Heliomicin-CsgA hydrogels further reduced fungal burden and suppressed hyphal development in an *ex vivo* porcine skin infection model. These findings demonstrate that microbial extracellular protein nanofibers can encode intrinsic biological functions that can be uncovered and further enhanced through protein engineering, expanding the design space for intrinsically bioactive antimicrobial biomaterials.

## Introduction

Microbially produced extracellular protein nanofibers have emerged as a versatile platform for genetically programmable biomaterials [1,2] By encoding material synthesis, self-assembly, and functionalization within microorganisms, these protein nanofibers have enabled a new generation of autogenic engineered living materials (ELMs) that integrate biological functions with material fabrication [3–7] Their modularity has facilitated the development of hydrogels, films, and three-dimensional architectures with tunable mechanical, chemical, and biological properties [8,9] Among microbial protein nanofibers, curli, composed of the major structural biofilm protein CsgA, has become one of the most extensively engineered autogenic ELM platforms due to its robust extracellular self-assembly, genetic accessibility, and modular architecture [10,11] Genetic fusion of bioactive peptides and proteins to CsgA has enabled diverse functions, including biotic and abiotic surface adhesion, bioremediation, catalysis, and tunable mechanical properties, while preserving nanofiber assembly [8,12,13] More recently, CsgA nanofibers have been engineered as therapeutic biomaterials displaying trefoil factors for the treatment of inflammatory bowel disease and viral antigens that function as self-adjuvanted vaccine platforms [8,10,13–16] Beyond the engineered fibers themselves, this therapeutic potential extends to biomanufactured hydrogels assembled directly from these fibers, which retain the bioactivity of their nanofiber building blocks. Previous work has shown that these engineered mucoadhesive nanofibers retain this functionality when assembled into hydrogels, enabling localized and prolonged therapeutic delivery to the gastrointestinal mucosa [8]

Despite extensive engineering of CsgA, its use as a scaffold for antifungal activity remains largely unexplored. Introducing antimicrobial function into autogenic ELMs would broaden their utility across biomedical applications, particularly against fungal pathogens such as *Candida albicans* (*C. albicans*), a leading cause of both mucosal and life-threatening systemic infections, particularly among immunocompromised and hospitalized patients [17,18] *C. albicans* frequently enters the body through disrupted skin or mucosal barriers, such as surgical sites, ulcers, or wounds, and accounts for nearly half of all cases of invasive candidiasis and candidemia, contributing to over 1.5 million infections and mortality exceeding 60% worldwide [18–20] Rising rates of antifungal resistance, combined with the limited number of available antifungal drug classes, have created an urgent need for new antifungal biomaterials capable of controlling fungal colonization and infection.

Here, we engineered CsgA nanofibers as a proof of concept for antimicrobial function against *Candida albicans* **(Figure 1)**. Antimicrobial peptides (AMPs) are an attractive class of antifungal agents. Many act through cationic interactions with negatively charged fungal membrane lipids, allowing for preferential selectivity, unlike broad-spectrum antifungal drugs, which disrupt the microbiome and promote secondary infection. [19–21] Because their primary mechanism involves membrane disruption rather than inhibition of a single molecular target, antifungal peptides are also less prone to the resistance mechanisms that undermine frontline antifungals, such as ERG11 mutations that drive azole resistance [22] Despite this promise, clinical translation of AMP-based therapies remains limited: (1) peptides are inherently unstable and rapidly degraded in biological environments, particularly at infection sites such as chronic wounds, where elevated protease activity and reactive oxygen species compromise stability and retention, and (2) production costs can reach hundreds of thousands of dollars per gram [23] Displaying genetically encoded antifungal peptides on a self-assembling, nanofiber scaffold offers a route to address these barriers by providing a stable, localized, and scalable delivery format. As a model antifungal peptide, we selected heliomicin, a 44-amino-acid defensin-family peptide produced by the tobacco budworm moth *Heliothis virescens* in response to infection. [21,24] Heliomicin adopts a cysteine-stabilized α-helix/β-sheet fold, a structural motif shared with plant and insect defensins and displays potent antifungal activity. [21,24]

**Figure 1.**
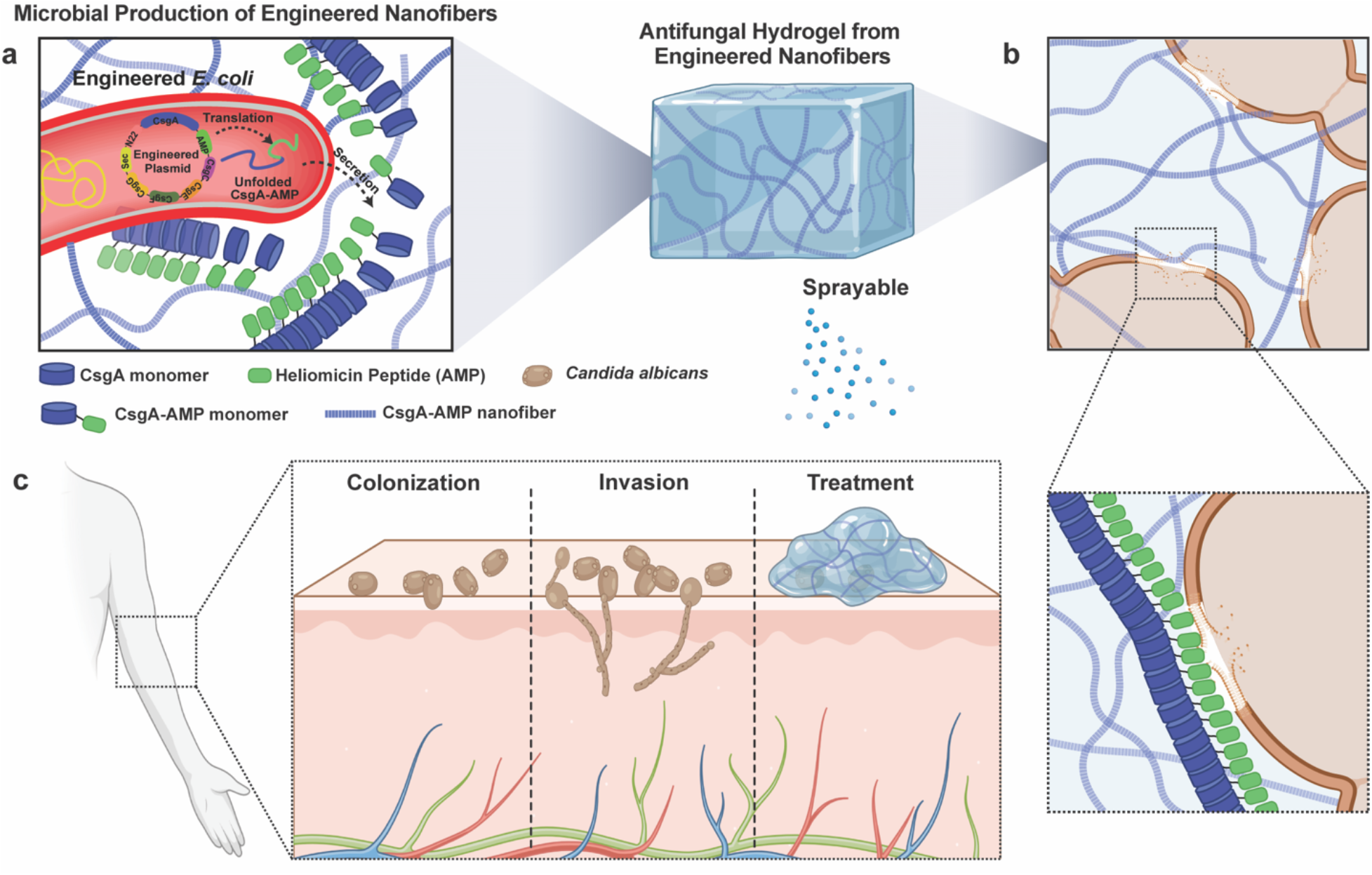
Engineering curli nanofibers for intrinsically active antifungal biomaterials. (a) Engineered *E. coli* produce and secrete Hel-CsgA monomers, which self-assemble into protein nanofibers and are utilized for biomanufacturing hydrogels. (b) Proposed mechanism of antifungal activity against C. albicans, highlighting interactions between the engineered nanofibers and the fungal cell membrane, (c) Potential application of the engineered hydrogel for localized treatment of topical fungal infection. Created in BioRender. Duraj-Thatte, A. (2026).

During evaluation of the engineered nanofibers, we unexpectedly discovered that wtCsgA nanofibers alone exhibit intrinsic antifungal activity against *C. albicans*, independent of heliomicin, revealing a previously unrecognized intrinsic biological activity. This observation suggests unreported anticandidal activity in the CsgA scaffold rather than serving solely as a passive structural scaffold for AMPs. Furthermore, genetic fusion of heliomicin significantly potentiated this intrinsic anticandidal activity, demonstrating that the intrinsic bioactivity of CsgA can be rationally enhanced through genetic engineering. More broadly, our results illuminate a new design principle for microbial protein nanofibers: engineered functions can be built on an inherently bioactive scaffold rather than a biologically inert structural framework, positioning CsgA nanofibers as a programmable platform for antimicrobial biomaterials.

## Results

### Genetic fusion of heliomicin preserves CsgA nanofiber self-assembly and hydrogel properties

To produce antifungal CsgA nanofibers, heliomicin was genetically fused to the N-terminus of CsgA through a flexible linker, downstream of the native Sec secretion signal and upstream of the N22 domain (N-terminal curli-specific targeting sequence) (**Figure SI1-2**). We selected this orientation to preserve the structural features required for heliomicin, as its N-terminal region is critical for antifungal activity. [21,24] The AlphaFold3 structural model of the fusion predicts that positioning heliomicin at the N-terminus maintains its functionally important N-terminal region in a solvent-exposed orientation while fusing its C-terminus to CsgA (**Figure 2b, Figure SI1**). The construct was expressed in the *Escherichia coli* (*E. coli*) PQN4 strain, which lacks the native curli operon and enables controlled production of engineered curli nanofibers (**Figure 2a, Figure SI2 & Table SI1**). The resulting Heliomicin-CsgA (Hel-CsgA) fusion protein was secreted and self-assembled extracellularly into curli nanofibers designed to display heliomicin on their surface. Scanning electron microscopy revealed dense fibrous networks secreted from cells that were morphologically indistinguishable from those produced by wild-type CsgA (wtCsgA) (**Figure 2c**), indicating that heliomicin is compatible with curli nanofiber assembly. Next, nanofiber production was quantified using Congo Red (CR), a dye that selectively binds amyloid nanofibers [25,26]. Both wtCsgA and Hel-CsgA *E. coli* strains exhibited significantly greater CR binding than the empty control (*E. coli* PQN4 lacking curli production), confirming robust extracellular nanofiber production in both strains. No significant difference in CR binding was observed between wtCsgA and Hel-CsgA, indicating that fusion of heliomicin does not impair curli nanofiber production (**Figure 2d**).

**Figure 2.**
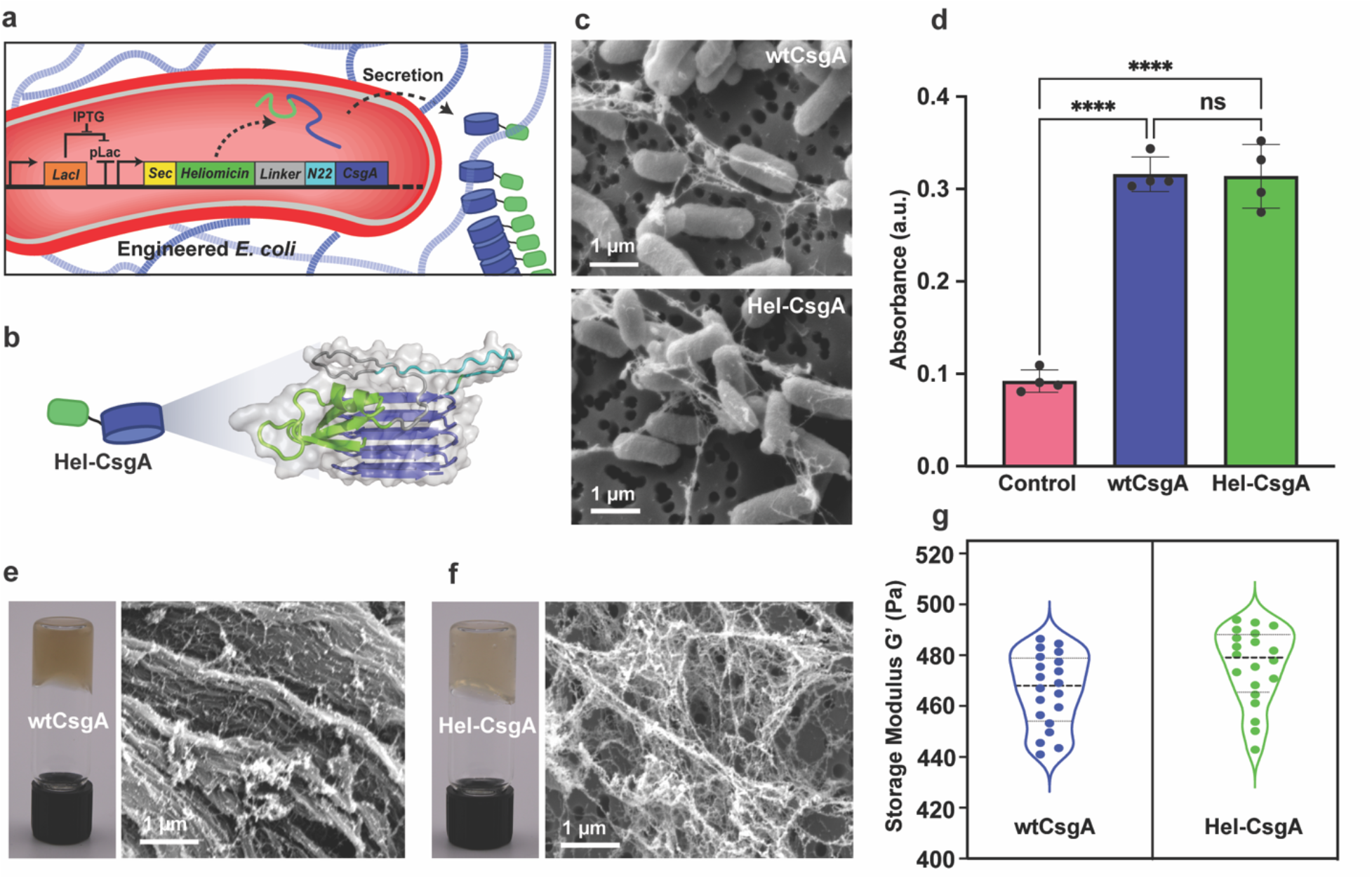
Characterization of engineered antifungal nanofibers and hydrogel. (a) Schematic of the genetic construct and microbial production of Hel-CsgA nanofibers, (b) AlphaFold3-predicted structure of the Hel-CsgA fusion protein, (c) SEM images of wtCsgA and Hel-CsgA nanofibers. Scale bars, 1 µm, (d) Congo Red binding assay evaluating the effect of heliomicin fusion on nanofiber production. Biological replicates n = 4. Data represented as mean ± standard deviation. ns non-significant, ****p≤0.0001. (e) Macroscopic and SEM imaging of wtCsgA hydrogel. Scale bar 1 µm. (f) Macroscopic and SEM imaging of Hel-CsgA hydrogel. Scale bars 1 µm. (g) Rheological analysis comparing the storage modulus (G’) of wtCsgA and Hel-CsgA hydrogels. Statistical significance was assessed by one-way ANOVA with Tukey’s post hoc test (*p<0.05). Parts of the schematic were created in BioRender. Duraj-Thatte, A. (2026).

Subsequently, to demonstrate the biomanufacturing of macroscopic biomaterials, hydrogels were fabricated directly from wtCsgA and Hel-CsgA producing nanofiber cultures using a filtration-based approach. [27] Both variants formed self-supporting macroscopic hydrogels. Scanning electron microscopy of the resulting hydrogels revealed dense, interwoven nanofiber networks that were morphologically similar, indicating that heliomicin fusion does not alter hydrogel microstructure (**Figure 2e-f, Figure SI3a-d**). Comparative rheological analysis showed that Hel-CsgA and wtCsgA hydrogels exhibited similar storage moduli (G’), ranging from approximately 432.85 ± 76.39 Pa for wtCsgA and 525.30 ± 121.32 Pa for Hel-CsgA, indicating that the heliomicin fusion did not significantly alter hydrogel mechanical stiffness (**Figure 2g**). [25] Together, these results demonstrate that genetically fusing heliomicin to CsgA preserves the scaffold’s native production, self-assembly, hydrogel to microstructural organization, and bulk mechanical properties.

### wtCsgA hydrogels exhibit antifungal activity enhanced by heliomicin fusion

The antifungal activity of Hel-CsgA hydrogels was evaluated by directly co-incubating *C. albicans* with Hel- or wtCsgA hydrogels for 48 h, followed by colony-forming unit (CFU) analysis (**Figure 3a-b, Figure SI4**). Unexpectedly, the wtCsgA hydrogel also reduced fungal viability by approximately 2 logs relative to untreated controls (**Figure 3b**), revealing a previously unrecognized intrinsic antifungal activity of the wtCsgA biomaterial. Hel-CsgA hydrogels produced a substantially greater reduction in fungal viability, decreasing CFU by approximately 3.5 logs relative to untreated controls and by an additional 1.5 logs compared with wtCsgA hydrogels (**Figure 3b**), demonstrating that genetic fusion of heliomicin significantly enhances the antifungal activity of the CsgA scaffold. Based on standard microbiological criteria, these results indicate a transition from fungistatic activity in wtCsgA hydrogels (<3-log reduction in CFU) to fungicidal activity in Hel-CsgA hydrogels (>3-log reduction in CFU). [28]

**Figure 3.**
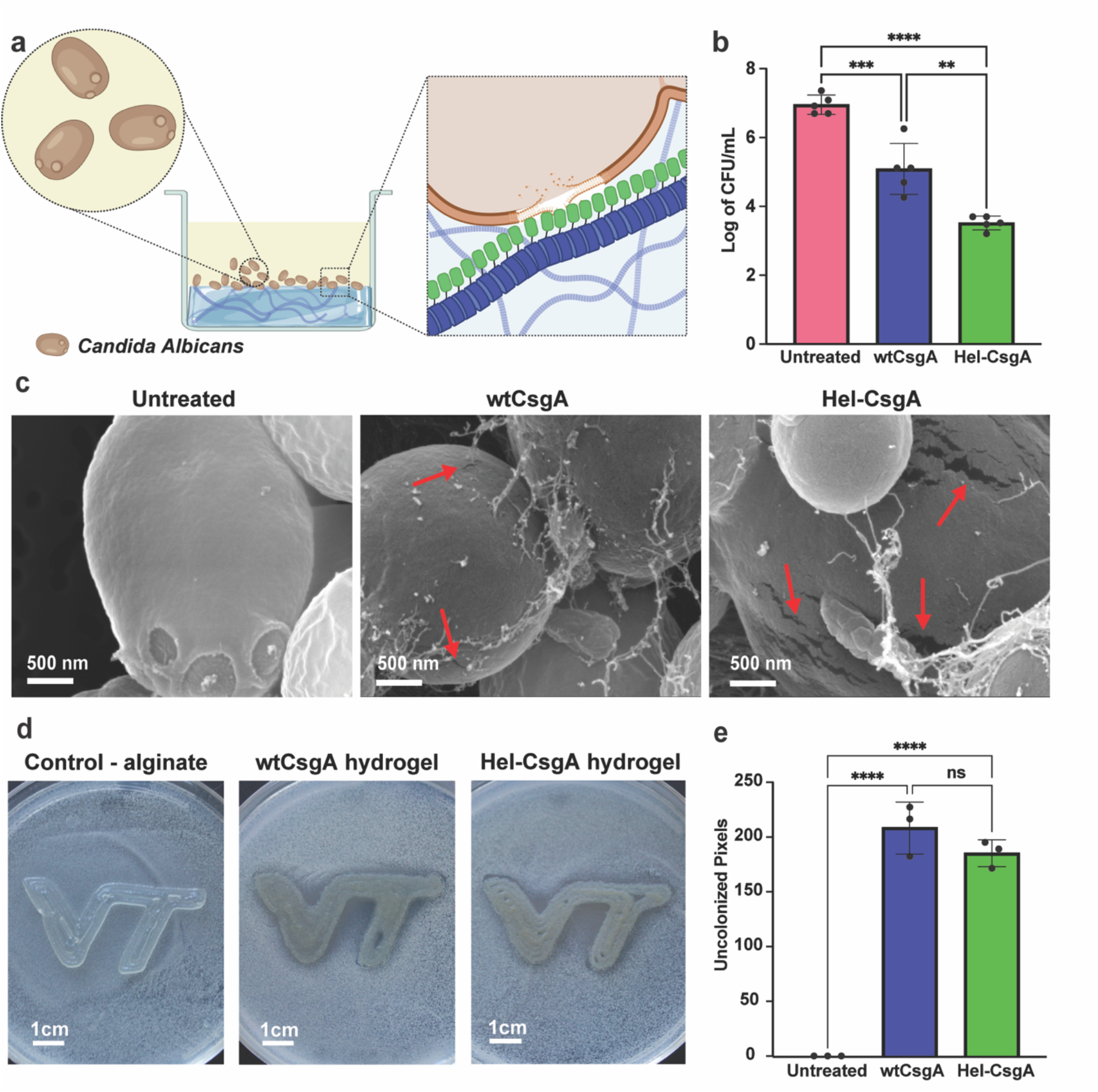
Intrinsic antifungal activity of CsgA nanofibers is enhanced by heliomicin fusion. (a) Schematic of the C. albicans challenge assay and proposed interaction between Hel-CsgA nanofibers and the fungal cell membrane, (b) Viable *C. albicans* quantified by colony-forming units (CFU) following 48 h incubation with wtCsgA or Hel-CsgA hydrogels, compared with the alginate control. Biological replicates n = 5. Data are presented as mean ± standard deviation. **p≤0.01, **p≤0.001, ***p≤0.0001. (c) SEM images of C. albicans cells following 48 h incubation without treatment or with wtCsgA or Hel-CsgA hydrogels. Red arrows indicate regions of fungal cell surface and membrane disruption. Scale bars, 500 nm, (d) Representative images of alginate control, wtCsgA, and Hel-CsgA hydrogels 3D printed onto C. albicans-colonized agar and incubated for 24 h at 30 °C. Scale bars, 1 cm, (e) Quantification of uncolonized areas surrounding the 3D-printed biomaterials using ImageJ. Biological replicates, n = 3. Data are presented as mean ± standard deviation. ns non-significant, ****p≤0.0001. Significance was assessed by one-way ANOVA with Tukey’s post hoc test (*p<0.05). Parts of the schematic were created in BioRender. Duraj-Thatte, A. (2026).

To investigate the structural basis of this enhanced antifungal activity, we examined *C. albicans* cells by scanning electron microscopy after treatment. Untreated cells exhibited smooth, intact surfaces with no evidence of membrane damage (**Figure 3c**), while cells incubated with alginate, a negative control, displayed a similar morphology (**Figure SI5**). Cells treated with wtCsgA hydrogels exhibited moderate surface disruption, including localized membrane tears and membrane deformation (**Figure 3c, Figure SI6a-d**), paralleling the CFU assay. In contrast, cells exposed to Hel-CsgA hydrogels exhibited extensive membrane damage characterized by widespread lysis, frequent surface ruptures, and leakage of intracellular contents (**Figure 3c**, **Figure SI7a-d**). These morphological changes match the reported membrane-disruptive mechanism of heliomicin and support the enhanced antifungal activity of Hel-CsgA nanofibers. [24]

To determine whether the antifungal activity was retained following 3D printing, alginate, wtCsgA, and Hel-CsgA hydrogels were printed directly onto *C. albicans* colonized agar plates (**Figure 3d**). In contrast to the alginate print, which became fully colonized by *C. albicans*, both Hel- and wtCsgA prints generated localized regions that remained free of fungal colonization, demonstrating that the antifungal activity was retained following 3D printing. Quantification of the uncolonized area from the printed constructs (**Figure 3e, Figure SI8a-c**) confirmed that both hydrogels produced significantly larger uncolonized regions than the alginate control, while no significant difference was observed between the two prints. The localized uncolonized regions are consistent with limited hydrogel spreading, suggesting that antifungal activity remained spatially localized to the printed hydrogels. These results demonstrate that the antifungal activities of wtCsgA and Hel-CsgA nanofiber biomaterials are preserved following hydrogel fabrication and 3D printing while remaining spatially localized to the printed constructs.

### Heliomicin fusion enhances membrane disruptive properties of CsgA biomaterials

To evaluate whether Hel-CsgA retained the disruptive features associated with heliomicin, we assessed *C. albicans* membrane integrity using two complementary fluorescence-based assays. Propidium iodide (PI), a membrane impermeant dye that fluoresces upon binding nucleic acids in cells with compromised membranes, was used to assess membrane permeabilization, with heat-killed cells serving as a positive control (**Figure SI9**). [29] Cells treated with wtCsgA biomaterials exhibited significantly greater PI fluorescence than untreated controls (**Figure 4a**), indicating increased membrane permeabilization in *C. albicans* treated with wtCsgA biomaterials. Hel-CsgA biomaterials produced a further increase in PI fluorescence compared with wtCsgA and untreated controls (**Figure 4a**), demonstrating enhanced membrane permeabilization following heliomicin fusion. Next, we further evaluated membrane disruption using the Nile Red assay, a lipophilic fluorescent dye that accumulates in intracellular membranes and is exported by ATP-binding cassette (ABC) transporters. [30] Cells treated with wtCsgA biomaterials exhibited a modest, non-significant increase in Nile Red release compared with untreated controls (**Figure 4b**), whereas Hel-CsgA biomaterials showed a significant increase in Nile Red release (**Figure 4b, Figure SI10**). The increased Nile Red release is consistent with membrane damage and supports the membrane-disruptive phenotype observed by SEM. The PI and Nile Red assays support membrane disruption by Hel- and wtCsgA, with substantially greater membrane-disruptive activity observed following heliomicin fusion.

**Figure 4.**
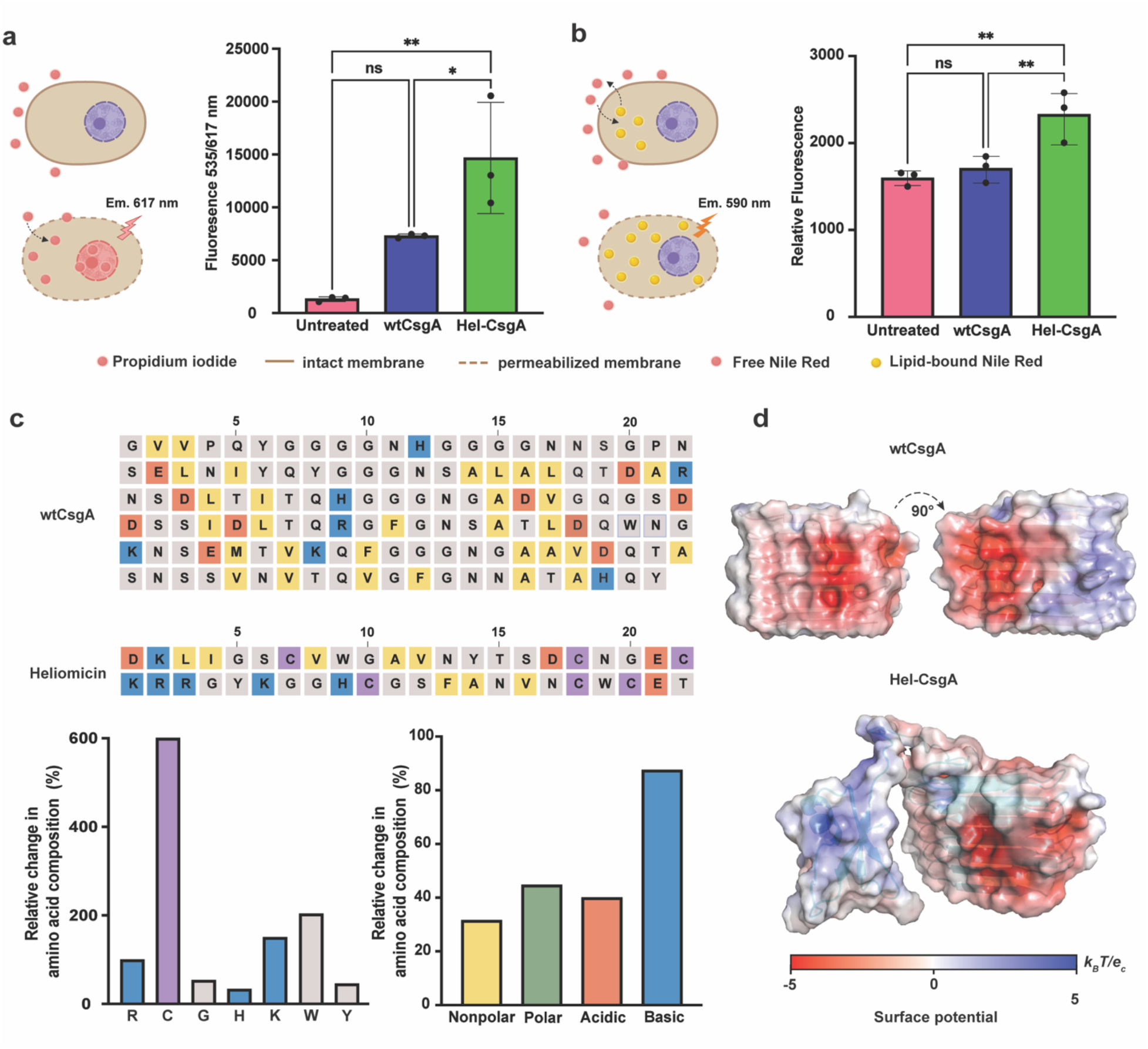
Heliomicin fusion enhances the membrane-disruptive properties and cationic surface of CsgA nanofibers. (a) Propidium iodide (PI) assay evaluating *C. albicans* membrane permeabilization following treatment with wtCsgA and Hel-CsgA hydrogels. Biological replicates, n = 3. Data are presented as mean ± standard deviation. *p≤0.0136, ****p≤0.0001, (b) Nile Red assay evaluating membrane-associated dye release from C. albicans following treatment with wtCsgA and Hel-CsgA hydrogels. Biological replicates, n = 3. Data are presented as mean ± standard deviation. ns non-significant, **p≤0.005, (c) Comparison of the primary sequences and amino acid composition of wtCsgA and heliomicin, highlighting differences in physicochemical residue classes, graphs below present, relative changes in amino acid composition following fusion of heliomicin to CsgA (left) and relative changes in the physicochemical composition of Hel-CsgA compared with wtCsgA (right). (d) Predicted electrostatic surface potentials of wtCsgA, heliomicin, and Hel-CsgA, illustrating changes in surface charge following heliomicin fusion. Schematic created in BioRender. Duraj-Thatte, A. (2026).

To investigate the molecular features associated with enhanced membrane-disrupting activity, we compared the primary sequence, amino acid composition, and predicted electrostatic surface properties of wtCsgA and Hel-CsgA. Sequence analysis revealed marked differences in amino acid composition between the two proteins, with heliomicin enriched in positively charged lysine and arginine residues, as well as cysteine residues characteristic of defensin peptides (**Figure 4c-d, Table SI2-4**). These sequence differences, along with amino acid analysis, demonstrated that Hel-CsgA exhibits an altered amino acid composition compared with wtCsgA, including increased proportions of nonpolar, polar, acidic, and basic residues, with the most pronounced increase observed for basic amino acids (**Figure 4e, Table SI3-4**). Electrostatic surface modeling revealed that wtCsgA possesses a localized positively charged surface patch, whereas heliomicin displays a substantially larger and more uniformly cationic surface (**Figure 4f**). The positively charged surface patch on CsgA is consistent with the membrane-permeabilizing activity observed experimentally and may contribute to its previously unrecognized antifungal activity. [21]

Together with the enrichment of basic amino acids in heliomicin, these features are consistent with the enhanced membrane permeabilization and antifungal activity observed following fusion to CsgA. Collectively, the sequence, electrostatic, PI, and Nile Red analyses suggest that expansion of the cationic surface contributes to the enhanced membrane disruptive activity and fungicidal behavior of Hel-CsgA.

### Hel-CsgA hydrogels reduce fungal burden and hyphal formation in an *ex vivo* porcine skin infection model

To evaluate the antifungal activity of Hel-CsgA hydrogels in a physiologically relevant tissue environment, an *ex vivo* porcine skin infection model adapted from Johnson *et al*., 2022 was established (**Figure 5a**). [31] Excised porcine skin explants were infected with *C. albicans* for 24 h to establish fungal colonization, followed by treatment with wtCsgA or Hel-CsgA hydrogels, or no treatment for an additional 24 h. Fungal colonization was evaluated by Grocott’s methenamine silver (GMS) staining, which selectively labels fungal cell walls, and quantified as the percentage of tissue area occupied by GMS-positive fungal structures. [32]

**Figure 5.**
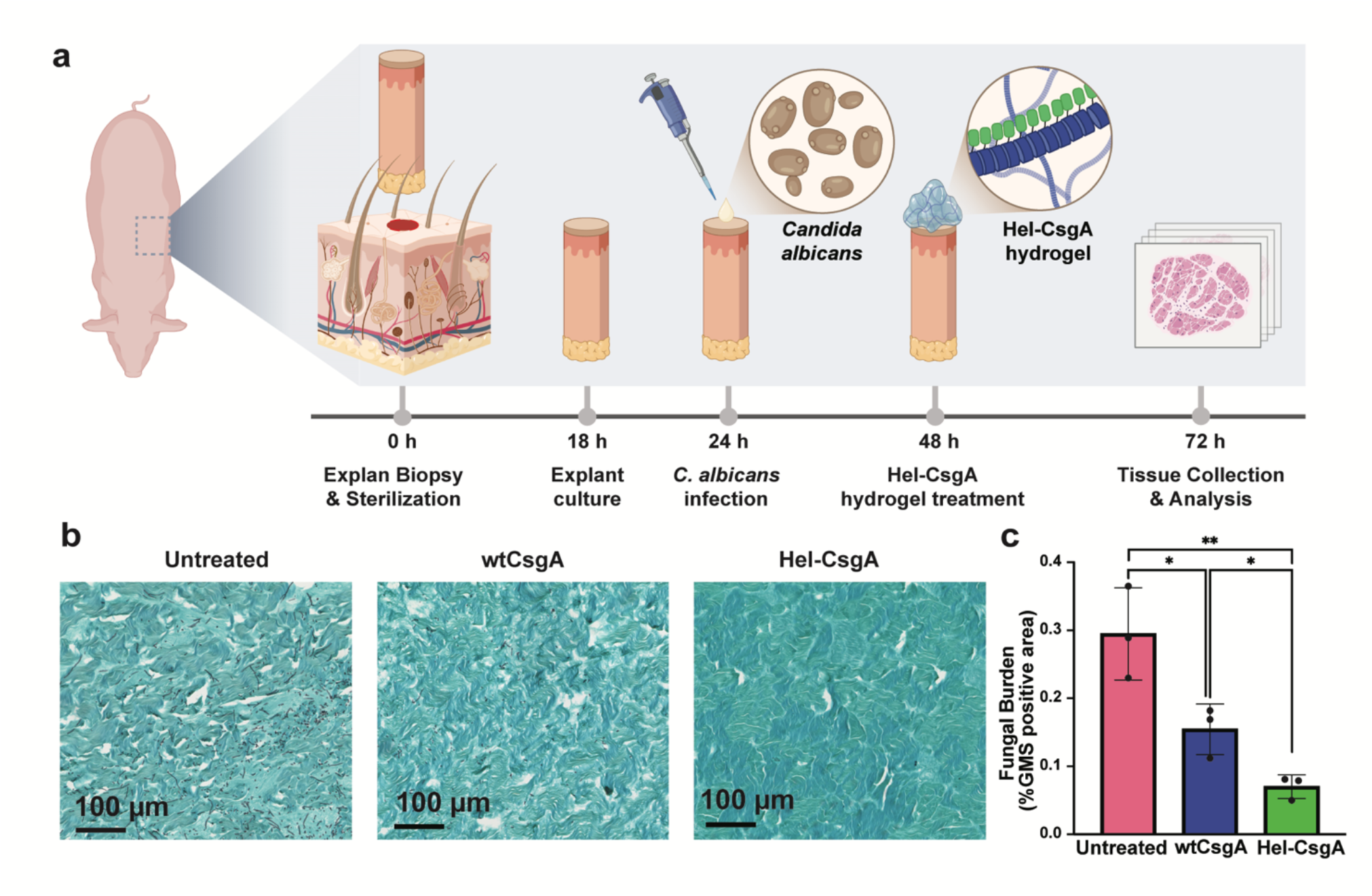
Antifungal activity of engineered CsgA hydrogels in an ex vivo C. albicans infection model. (a) Schematic and experimental timeline of the ex vivo porcine skin infection model and hydrogel application, (b) Representative GMS-stained histological images of C. albicans-infected skin explants following no treatment or treatment with wtCsgA or Hel-CsgA hydrogels. Scale bars, 100 µm, (c) Quantification of GMS-positive fungal area using Trainable Weka Segmentation in ImageJ. Biological replicates, n = 3. Data represented as mean ± standard deviation. *p≤0.035, **p≤0.0051. Significance was assessed by t-Test (*p<0.05). Schematic created in BioRender. Duraj-Thatte, A. (2026).

Untreated explants showed extensive fungal colonization, with abundant yeast and hyphal structures distributed throughout the tissue (**Figure 5b, Figure SI11a-e**). Treatment with wtCsgA hydrogels significantly reduced fungal burden compared with untreated controls (**Figure 5b,c, Figure SI12a-e**). Hel-CsgA hydrogels produced a further reduction in fungal burden, decreasing the GMS-positive fungal area by approximately 76% relative to untreated tissue and by approximately 55% relative to wtCsgA hydrogels (**Figure 5c, Figure SI13a-e**). Histological examination further revealed differences in fungal morphology following treatment. While untreated explants contained abundant hyphal structures penetrating the tissue, wtCsgA-treated explants exhibited reduced fungal colonization but retained visible hyphae (**Figure 5b**). In contrast, Hel-CsgA-treated explants displayed markedly fewer hyphal structures, with fungal cells observed predominantly in the yeast form (**Figure 5b**). These observations demonstrate that Hel-CsgA hydrogels reduce fungal burden and are associated with a marked reduction in hyphal structures in infected tissue.

## Discussion

Microbial extracellular protein nanofibers have emerged as a versatile platform for ELMs, as they can be genetically modified to display diverse biological functions while preserving their ability to self-assemble into extracellular materials. [12,33] In this study, we genetically engineered CsgA nanofibers to display the antifungal peptide heliomicin and demonstrated that genetic fusion significantly enhances antifungal activity while preserving nanofiber self-assembly, hydrogel formation, and 3D printability. Evaluation of the engineered nanofibers revealed that wtCsgA itself possesses intrinsic antifungal activity against *C. albicans*, a biological function that, to our knowledge, has not been reported previously. These findings reveal that CsgA is not merely a structural scaffold for displaying functional peptides but rather an intrinsically bioactive protein-based biomaterial whose biological activity can be further enhanced through protein engineering. Furthermore, curli and various nanofibers have been extensively investigated as structural extracellular scaffolds implicated in bacterial adhesion, biofilm formation, surface colonization, and cellular protection. [15,34–36] Previous engineering strategies have largely focused on displaying functional peptides or proteins to introduce new biological functions, while the biological activity has generally been attributed to the displayed cargo rather than the scaffold itself. [8,36] Our findings demonstrate that CsgA itself possesses intrinsic biological activity, suggesting that extracellular protein nanofibers harbor previously unrecognized functions beyond structural support. [36–39]

The molecular basis of this intrinsic antifungal activity remains to be fully understood. However, our computational and experimental analyses provide several important insights. WtCsgA nanofibers increased membrane permeabilization, as demonstrated by PI uptake, and exhibited a localized positively charged surface region predicted by electrostatic modeling. Fusion of heliomicin increased the abundance of basic amino acids and expanded the positively charged surface of the nanofibers, changes that were associated with enhanced membrane disruption and a transition from fungistatic to fungicidal activity. [40] Given the extensive membrane damage also observed by scanning electron microscopy, electrostatic interactions between the positively charged nanofiber surface and the negatively charged fungal membrane likely contribute to this antifungal activity. Although future mutational studies will be required to directly determine the contribution of specific residues, the agreement between the computational predictions and experimental observations identifies surface charge as an important molecular feature associated with membrane-disruptive activity.

Importantly, the antifungal activity of both wtCsgA and Hel-CsgA was retained in an *ex vivo* porcine skin infection model, demonstrating that both biomaterials remain active in a tissue-relevant environment. Hel-CsgA biomaterials substantially reduced fungal burden and were associated with a marked reduction in hyphal structure compared with untreated tissue and wtCsgA-treated explants. As hyphal growth is a key virulence trait that promotes tissue invasion and persistence during *Candida* infections, these findings suggest that genetically engineered curli biomaterials may provide a platform for localized therapies. [41,42] More broadly, the retention of antifungal activity in precolonized *in vitro* and *ex vivo* infection models demonstrates the robustness of this biomaterial platform and its capacity for further functional enhancement through genetic peptide fusions. This work highlights the potential of engineered protein nanofibers as biomanufactured therapeutic materials.

Our findings highlight a novel strategy for engineering antimicrobial protein biomaterials. To date, researchers have primarily developed engineered therapeutic protein nanofibers by genetically displaying functional peptides or proteins on extracellular scaffolds, with biological activity largely attributed to the engineered cargo rather than the scaffold itself. In contrast, our results demonstrate that the scaffold itself can possess intrinsic biological activity that is further enhanced through protein engineering. Rather than introducing an entirely new biological function, heliomicin amplifies an existing CsgA property. These findings suggest that intrinsically bioactive biomaterials can be engineered by building upon biological functions encoded within extracellular protein nanofibers. Rather than relying on post-fabrication incorporation or release of antimicrobial agents, this platform encodes antimicrobial activity directly within the protein material itself. This approach offers new opportunities to engineer pathogen-specific biomaterials by combining the intrinsic activities of natural protein scaffolds with genetically encoded functional domains.

The evolutionary origin of the intrinsic antifungal activity of CsgA remains unknown. Curli nanofibers are major structural components of bacterial biofilms, which often form complex polymicrobial communities comprising bacteria, fungi, and other microorganisms. [43,44] Within these polymicrobial communities, bacteria and fungi continuously interact and compete for nutrients, space, and ecological niches. [45] We therefore hypothesize that the intrinsic antifungal activity of CsgA can represent an evolutionarily conserved ecological function that contributes to regulating fungal colonization within mixed-species biofilms. In this context, the localized positively charged surface identified in wtCsgA may have evolved not only to support biofilm architecture but also to mediate interactions with neighboring fungal cells. Although this hypothesis requires experimental validation, it suggests that extracellular protein nanofibers may play a broader role in microbial interactions than currently described.

### Summary

Collectively, our findings identify a previously unrecognized intrinsic antifungal activity of CsgA nanofibers and demonstrate that this activity can be substantially enhanced through protein engineering. More broadly, this work suggests that microbial extracellular protein nanofibers should not be viewed solely as structural scaffolds, but rather as naturally bioactive protein materials whose intrinsic functions can be discovered and systematically engineered. This study proposes a new platform for developing intrinsically bioactive biomaterials, in which antimicrobial functions are encoded directly within the material itself and further enhanced through rational protein engineering. We anticipate that extending this concept to the rapidly expanding repertoire of naturally occurring extracellular protein nanofibers (β-solenoid proteins) recently reported will facilitate both the discovery of new biological functions and the development of programmable biomaterials with tailored therapeutic activities. [36,37]

## Author Contributions

A.D.T., A.K., and N.B. developed the initial concept. A.D.T., A.K., and N.B. streamlined the central concept and methodology. A.D.T. and N.B. designed experiments. A.K. performed initial experiments on engineered antifungal nanofibers and hydrogels. N.B. performed colony-forming unit plating, propidium iodide and Nile Red assays, *ex vivo* porcine model, and scanning electron microscopy imaging of nanofibers, hydrogels, and their characterization. A.K. and H.H designed and engineered Hel-CsgA nanofibers. B.R. and M.B. performed molecular cloning experiments. A.D.T. supervised N.B., A.K., H.H., B.R., and M.B. The manuscript was drafted by N. B., A. K., and A.D.T. with input from all authors. All authors discussed the results and commented on the manuscript.

## Acknowledgements

Work was partly performed at the BioMaterial Characterization Laboratory (BioMCL), Nanoscale Characterization and Fabrication Laboratory (NCFL), and Virginia Tech Animal Laboratory Services (ViTALs). This work was supported by the U.S. National Science Foundation EFRI-ELiS program under Award No. 2318093.

We thank Katherine Scott for training on the Weka system for fungal identification and quantification in histological images.

## Conflicts of Interest

A.D.T., N.B., and A.K. are inventors on a patent application related to the antimicrobial peptide engineering of the curli nanofiber scaffold and de novo autogenic engineered living materials described in this manuscript. “Programmable Bioactive Protein Nanofibers for Antimicrobial Applications”, U.S. Patent App.63/898,598

## STAR METHODS

- KEY RESOURCES TABLE
- EXPERIMENTAL MODEL AND ORGANISM DETAILS

- Bacterial strains, medium composition, and culture conditions
- METHOD DETAILS

- Fiber culture and harvest
- Congo Red dye binding assay
- Optical Images
- Hydrogel Rheology
- *C. albicans* biomaterial challenge and CFU
- SEM Imaging
- Propidium Iodide Staining
- Nile Red Release Assay
- 3D Print *C. albicans* challenge
- *Ex vivo* porcine model
- QUANTIFICATION AND STATISTICAL ANALYSIS

- Image Analysis
- Statistical Analysis
- RESOURCE AVAILABILITY

### Lead Contact

Further information and requests for resources should be directed to and will be fulfilled by the lead contact, Anna Duraj-Thatte

### Materials Availability

Plasmids generated in this study are available from the lead contact upon request.

### Data and Availability

All data supporting the findings of this study are available within the article and its supplementary information.

### KEY RESOURCES TABLE

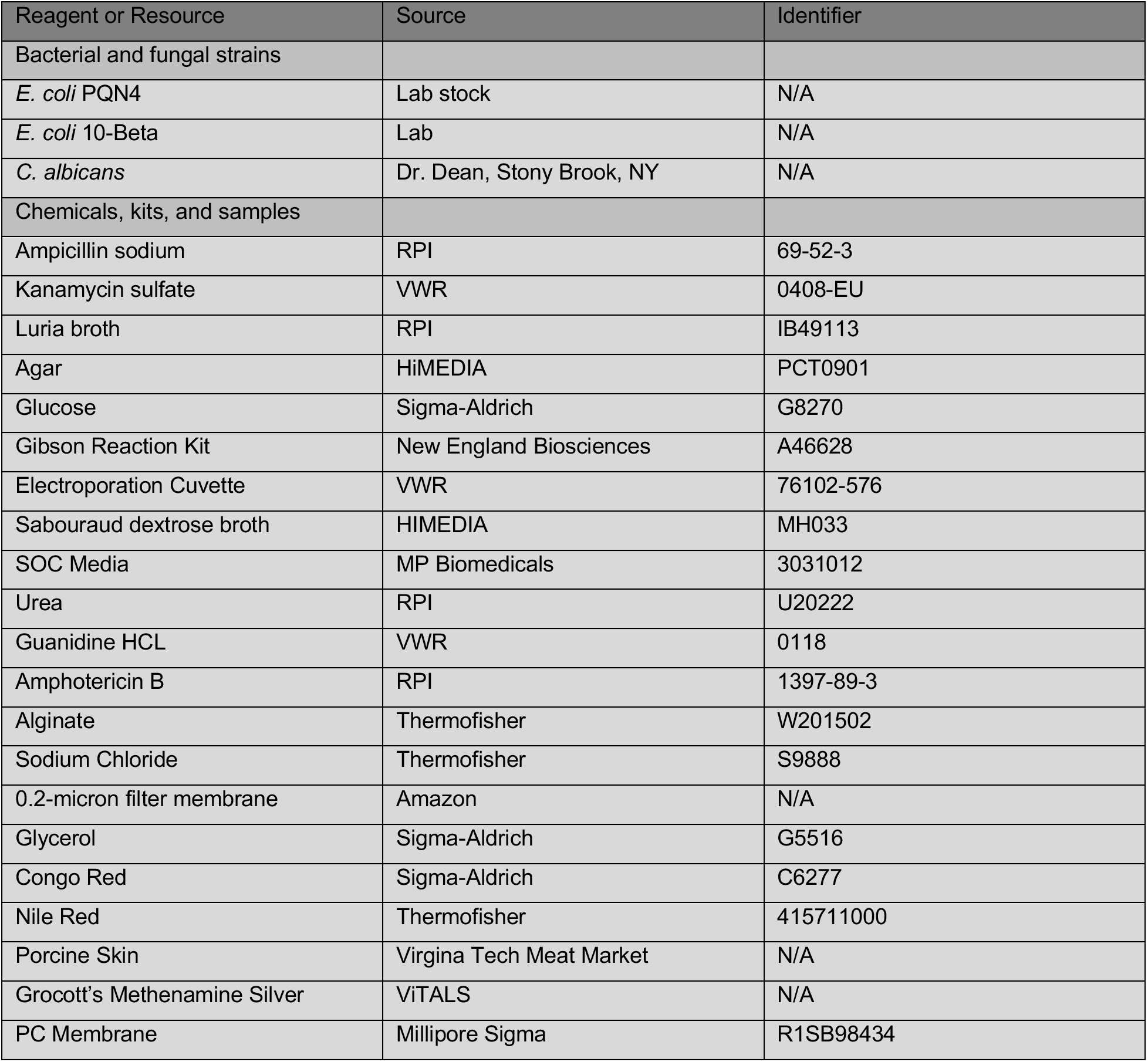
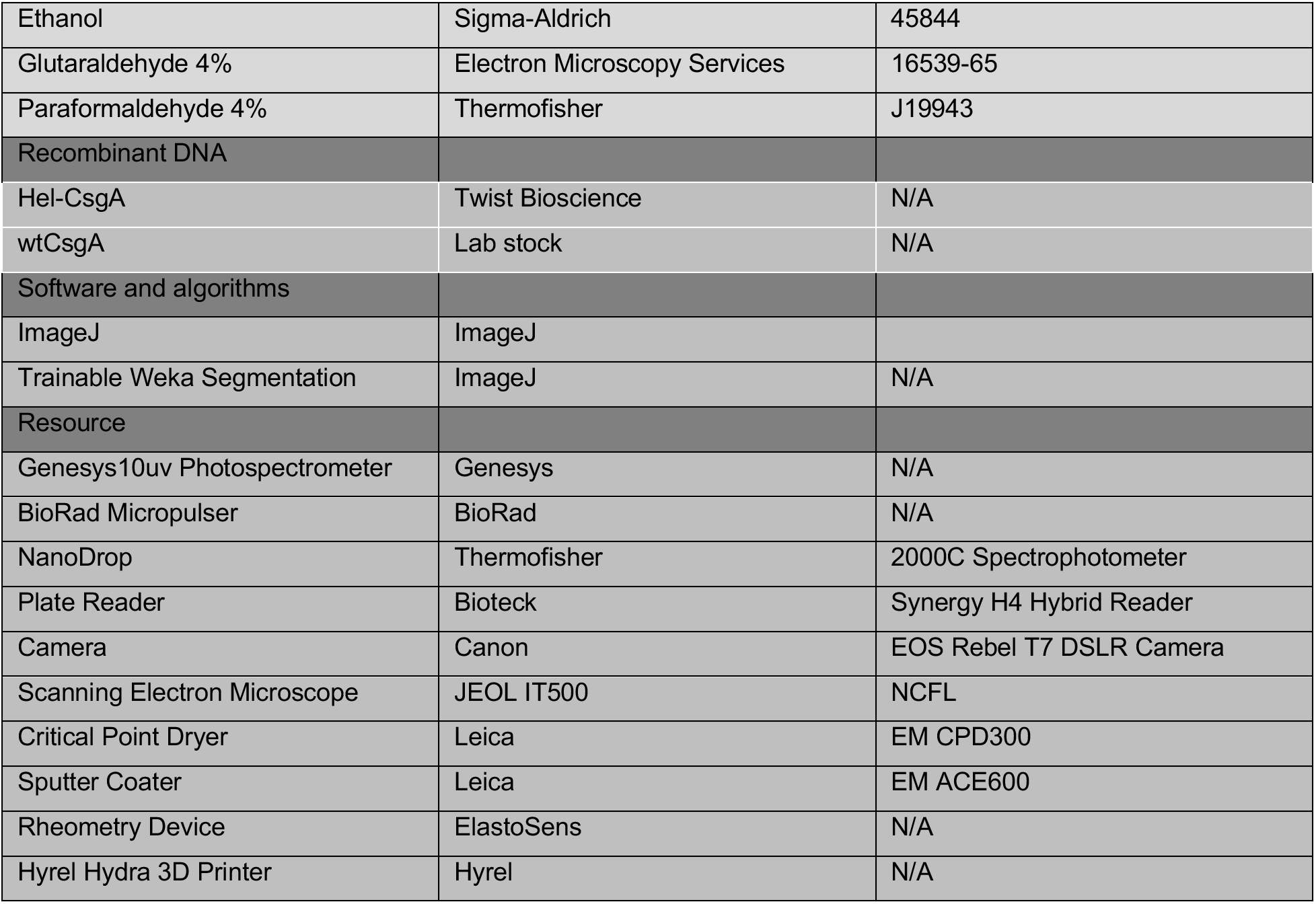

### EXPERIMENTAL MODEL AND SUBJECT DETAILS

#### Bacterial Strains and Culture Conditions

Wild-type CsgA (wtCsgA) and Heliomicin-CsgA (Hel-CsgA) plasmid constructs were generated in the pET21d vector using Gibson assembly. We confirmed correct assembly by whole-plasmid sequencing after transformation into *E. coli* 10-Beta. The curli operon knockout strain *E.coli* PQN4 was provided by Joshi lab *at Northeastern* University. Sequence-verified plasmids were subsequently transformed into *E. coli* PQN4 and stored at −80 °C in LB medium supplemented with 20% glycerol.

For nanofiber production, individual colonies were inoculated from freshly streaked agar plates containing ampicillin (100 µg mL⁻¹) and glucose (0.5%) and cultured overnight at 37 °C with shaking at 200 rpm. One milliliter of overnight culture was transferred into 500 mL of LB medium supplemented with ampicillin (100 µg mL⁻¹) and incubated for 48 h at 37 °C with shaking at 250 rpm.

#### Fungal Strains and Culture Conditions

*Candida albicans* was cultured overnight in Sabouraud dextrose broth (SDB) supplemented with kanamycin at 30 ^°^C and 200 rpm. Cultures were diluted to 5 x 10^7^ CFU mL^-1^ prior to antifungal assays.

### METHOD DETAILS

#### Production of wtCsgA and Hel-CsgA Hydrogels

Following 48 h of culture, bacterial suspensions were incubated for 45 min with 8 M urea at 4 °C and vacuum-filtered through 2 µm membranes. The retained material was subsequently treated with 8 M guanidine hydrochloride for 15 min, followed by washing and treatment with 5% SDS. The material was then extensively washed with distilled water and collected from the filter membrane to yield curli hydrogels. Hydrogels were stored at 4 °C until use.

#### Congo Red Binding Assay

Curli nanofiber production was quantified using Congo Red (CR) binding. Briefly, 1 mL of 48 h bacterial culture was centrifuged at 4,000 rpm for 10 min. Pellets were resuspended in 0.25 mM CR solution and incubated for 10 min at room temperature. Samples were centrifuged at 14,000 rpm for 10 min and supernatants were transferred to 96-well plates. Absorbance was measured using a BioTek Synergy H4 Hybrid Plate Reader at 414 nm. Measurements were performed using biological replicates (n =4) with five technical replicates per sample.

#### Hydrogel Rheology

Mechanical properties of wtCsgA and Hel-CsgA hydrogels were evaluated using an ElastoSens^TM^ Bio rheometer (Rheolution, Montreal, Canada). Approximately 300 mg of hydrogel was loaded into the sample holder and monitored for 2 min with measurements collected every 2 s. Data were collected from biological replicates (n = 3).

#### Antifungal Colony Forming Unit Assay

Antifungal activity was evaluated by co-incubating 100 µL of *C. albicans* suspensions (5 x 10^7^ CFU mL^-1^) with 100 mg of wtCsgA hydrogel, Hel-CsgA hydrogel, alginate hydrogel, amphotericin B loaded alginate hydrogel (1µg mL^-1^), or untreated controls in 96-well plates. Samples were incubated for 48 h at 30 ^°^C.

Following incubation, serial dilutions were prepared and plated onto SDB agar supplemented with kanamycin. Plates were incubated at 30 ^°^C overnight before colony counting. Experiments were performed using biological replicates (n = 5).

#### Scanning Electron Microscopy

To image fungal morphology, samples from antifungal assays were deposited onto 0.2 µm PC membrane filters and fixed for 4 hours at 4 ^°^C in a 1:1 solution of 4% paraformaldehyde and glutaraldehyde.

Samples were dehydrated through ethanol replacement solutions (25% to 100%), followed by critical point drying using a Leica EM CPD3000 system. Dried samples were sputter-coated with platinum using a Leica ACE600 sputter coater to a height of 18 nm. Images were then taken at 5kV and 50V probe current using a JEOL IT500 scanning electron microscope (NCFL, Virginia Tech).

#### Propidium Iodide Membrane Permeabilization Assay

Membrane permeabilization was evaluated using propidium iodide (PI). *C. albicans* cultures (100 µL, 5 x 10^7^ CFU mL^-1^) were incubated with wtCsgA hydrogels, Hel-CsgA hydrogels, or untreated controls for 48 h at 30 ^°^C. Heat-killed cells were generated by boiling at 100 ^°^C for 5 min and served as positive controls.

Following treatment, PI was added to a final concentration of 1 µg mL^-1^ and samples were incubated for 30 min at room temperature in the dark. Fluorescence was measured using a BioTek Synergy H4 Hybrid Plate Reader (Ex/EM = 535/617 nm). Measurements were performed using biological replicates (n = 3).

#### Nile Red Release Assay

Membrane disruption was further evaluated using Nile Red. Following 48 h incubation with hydrogels, fungal suspensions were stained with Nile Red (7 µM) for 20 min. Cells were pelleted by centrifugation and supernatants were transferred to 96-well plates. Fluorescence was measured using a BioTek Synergy H4 Hybrid Plate Reader (Ex/EM = 488/590 nm). Measurements were performed using biological replicates (n = 3).

#### Three-Dimensional Printing and Antifungal Challenge

Hydrogels were loaded into 5 mL BD syringes fitted with a 14-gauge nozzle and printed using a Hyrel 3D printed. A Virginia Tech logo (copyright free) was used as the printing template.

Hydrogels were printed directly onto SDB agar supplemented with kanamycin and seeded with a 1:100 dilution of an OD_600_ = 1 *C. albicans* culture. Following incubation for 24-48 h at 30 ^°^C, regions lacking fungal growth surrounding the printed structures were imaged and quantified. Experiments were performed using biological replicates (n = 3).

#### *Ex Vivo* Porcine Skin Infection Model

An *ex vivo* porcine skin infection model was adapted from Johnson C.J. et al (2022) with minor modifications. Porcine skin explants were placed in 24-well plates and inoculated with *C. albicans* (100 µL, 5 x 10^7^ CFU mL^-1^). Following 24 h colonization, explants were treated with wtCsgA or Hel-CsgA hydrogels or left untreated.

After an additional 24 h incubation, samples were fixed overnight in 70% ethanol at 4 ^°^C. They were then provided to ViTALs (Virginia Tech Animal Laboratory Services) for histology preparation and stained using Grocott’s methenamine silver staining for fungal visualization.

#### Optical Imaging

Macroscopic photographs of hydrogels and printed constructs were acquired using a Canon EOS Rebel T7 DSLR camera.

### QUANTIFICATION AND STATISTICAL ANALYSIS

#### Histological Image Analysis

GMS-stained tissue sections were imaged at 20× magnification. Images were analyzed using Fiji/ImageJ. ND2 image stacks were imported, and the central focal plane was selected for analysis. Fungal structures were segmented using the Trainable Weka Segmentation plugin trained to distinguish fungal cells from porcine tissue.

Fungal burden was quantified as the percentage of tissue area occupied by GMS-positive fungal structures. Measurements were obtained from duplicate images for each of the biological replicates (n = 3) and averaged prior to statistical analysis.

#### Statistical Analysis

All analysis was performed in GraphPad Prism version 10.2.0

Congo Red analysis of biological replicates (n = 3) was performed using One-way

ANOVA followed by Tukeys multiple comparison test. Resulting in wtCsgA to Hel-CsgA ns, Hel- or wtCsgA to Empty (ΔCsgA PQN4) p < 0.0001.

Colony forming unit assay was analyzed using biological replicates (n = 3) in a One-way ANOVA followed by Tukeys multiple comparison test. Resulting in untreated control to wtCsgA p = 0.0003, untreated control to Hel-CsgA p < 0.0001, wtCsgA to Hel-CsgA p = 0.002.

Analysis of 3D printed biomaterial clear zones was performed utilizing pixel counts from ImageJ software using biological replicates (n = 3) and analyzed using a One-way ANOVA followed by Tukeys multiple comparison test. Resulting in algiante control to wtCsgA p < 0.0001, alginate control to Hel-CsgA p < 0.0001, wtCsgA to Hel-CsgA p = 0.22.

Propidium Iodide staining analysis of biological replicates (n = 3) was performed using One-way ANOVA followed by Tukeys multiple comparison test. Resulting in untreated control to wtCsgA p < 0.0001, untreated control to Hel-CsgA p < 0.0001, wtCsgA to Hel-CsgA p = 0.014.

Nile Red release analysis of biological replicates (n = 3) was performed using a One-way ANOVA. followed by Tukeys multiple comparison test. Resulting in untreated control to wtCsgA p = 0.89, untreated control to Hel-CsgA p = 0.0012, wtCsgA to Hel-CsgA p = 0.004.

*Ex vivo* analysis of biological replicates (n = 3) was performed using a unpaired t-test. Resulting in untreated control to wtCsgA p = 0.035, untreated control to Hel-CsgA p = 0.0051, wtCsgA to Hel-CsgA p = 0.024.

## Notes

### Competing Interest Statement

The authors have declared no competing interest.

