## Supplementary file for "Intrinsic antifungal activity of curli nanofibers expands the design space for programmable antimicrobial biomaterials"

* Co-authors

### Corresponding author

**Supplementary Figure 1. Antifungal engineered Heliomicin-CsgA structure prediction using AlphaFold2.** The protein structure of CsgA, with N-terminal fusion of Heliomicin predicted by AlphaFold2, with plDDT (predicted local distance difference test) score markings.

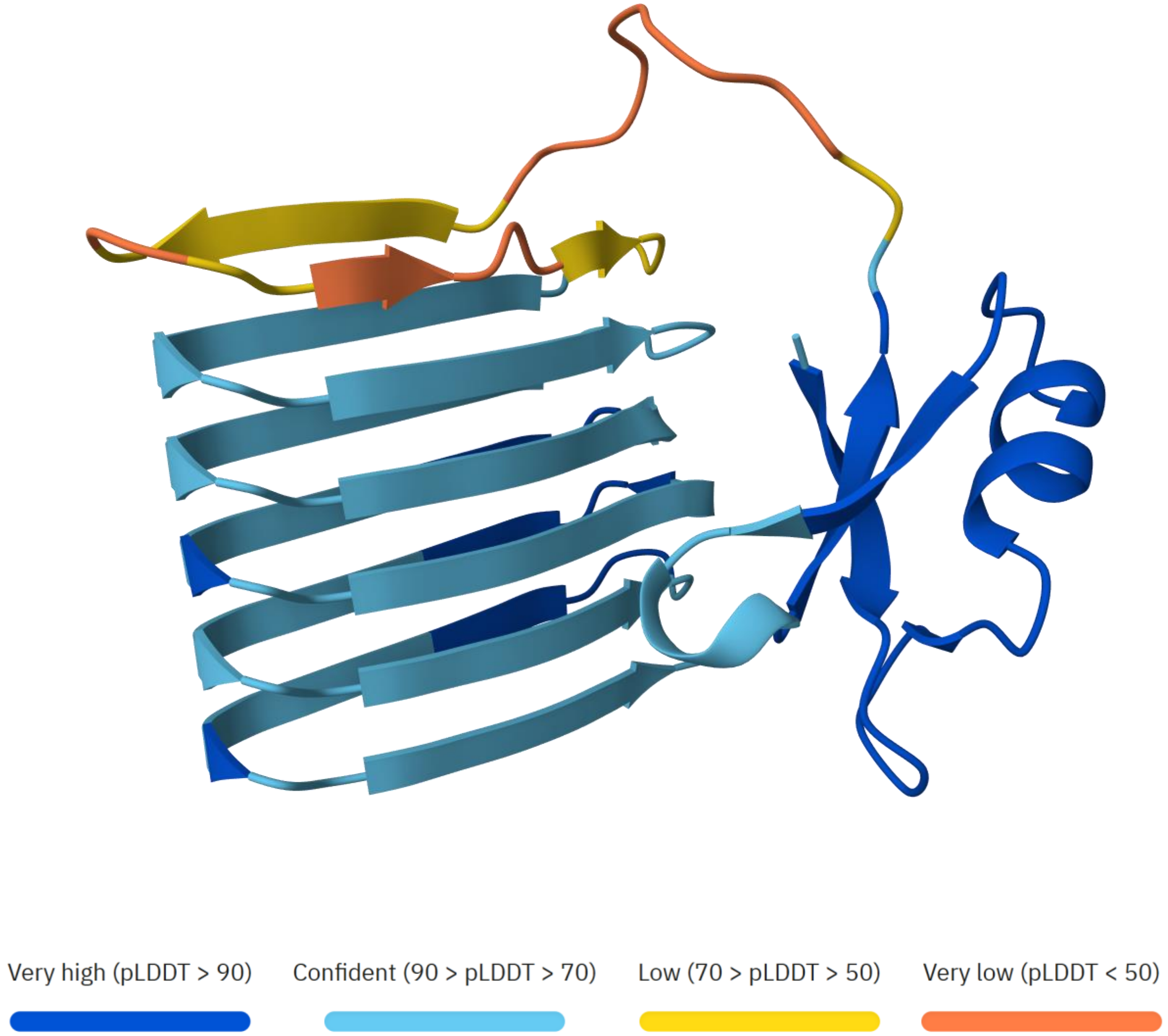

**Supplementary Figure 2. Plasmid map of antifungal engineered nanofibers**

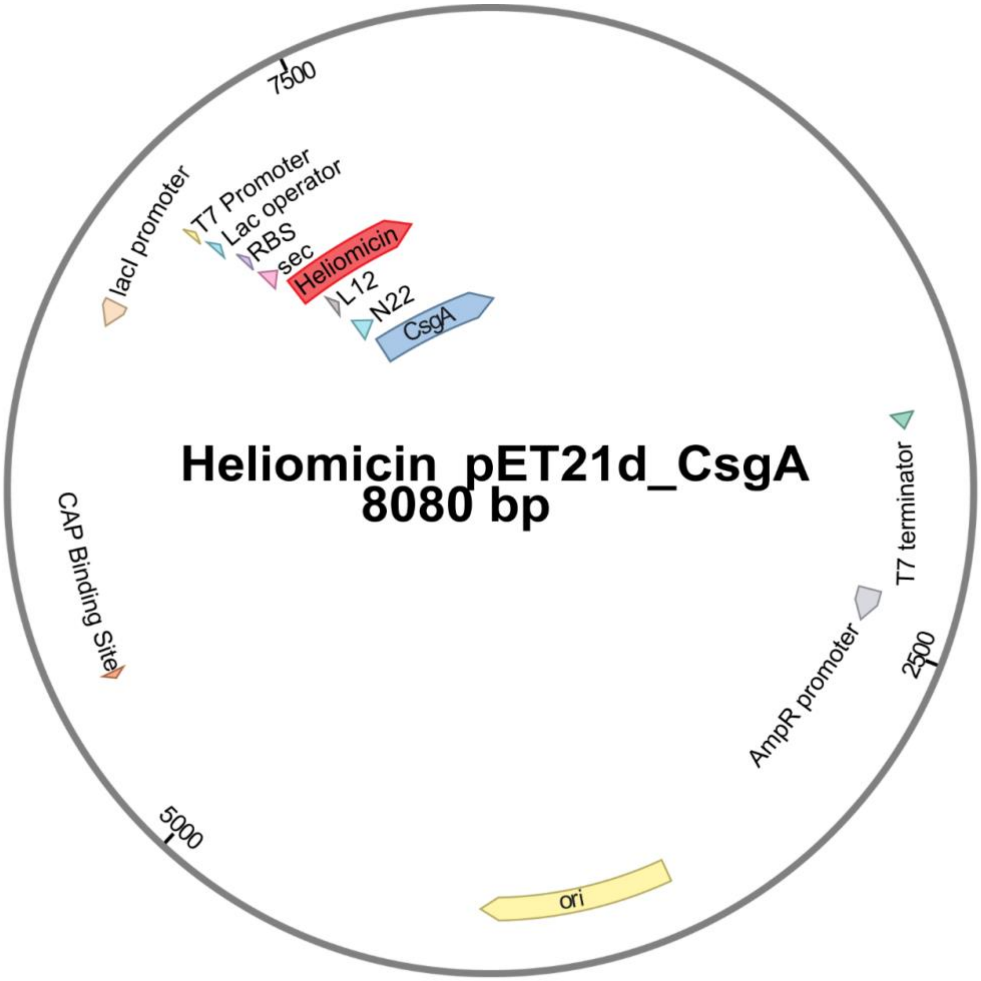

**Supplementary Table 1. Sequences of CsgA homologs and variants used in this study.**

| Protein | Protein Sequence | DNA Sequence |
|---|---|---|
| **CsgA** | GVVPQYGGGGNHGGGGNNSGPNSELNIYQYGG<br>GNSALALQTDARNSDLTITQHGGGNGADVGQGSD<br>DSSIDLTQRGFGNSATLDQWNGKNSEMTVKQFG<br>GGNGAAVDQTASNSSVNVTQVGFGNNATAHQY* | GGTGTTGTTCCTCAGTACGGCGGCGGCGGTAACCAC<br>GGTGGTGGCGGTAATAATAGCGGCCCAAATTCTGAG<br>CTGAACATTTACCAGTACGGTGGCGGTAACTCTGCA<br>CTTGCTCTGCAAACTGATGCCCGTAACTCTGACTTGA<br>CTATTACCCAGCATGGCGGCGGTAATGGTGCAGATG<br>TTGGTCAGGGCTCAGATGACAGCTCAATCGATCTGA<br>CCCAACGTGGCTTCGGTAACAGCGCTACTCTTGATC<br>AGTGGAACGGCAAAAATTCTGAAATGACGGTTAAACA<br>GTTCGGTGGTGGCAACGGTGCTGCAGTTGACCAGAC<br>TGCATCTAACTCCTCCGTCAACGTGACTCAGGTTGG<br>CTTTGGTAACAACGCGACCGCTCATCAGTACTAA |
| **Hel-CsgA** | DKLIGSCVWGAVNYTSDCNGECKRRGYKGGHCG<br>SFANVNCWCETGSGGSGGSGGSGGVVPQYGGG<br>GNHGGGGNNSGPNSELNIYQYGGGNSALALQTD<br>ARNSDLTITQHGGGNGADVGQGSDDSSIDLTQRG<br>FGNSATLDQWNGKNSEMTVKQFGGGNGAAVDQT<br>ASNSSVNVTQVGFGNNATAHQY* | GATAAACTGATTGGCAGCTGCGTGTGGGGCGCGGT<br>GAACTATACCAGCGATTGCAACGGCGAATGCAAACG<br>CCGCGGCTATAAAGGCGGCCATTGCGGCAGCTTTGC<br>GAACGTGAACTGCTGGTGCGAAACCGGGAGCGGCG<br>GCTCTGGCGGCTCTGGCGGTTCTGGCGGTGTTGTTC<br>CTCAGTACGGCGGCGGCGGTAACCACGGTGGTGGC<br>GGTAATAATAGCGGCCCAAATTCTGAGCTGAACATTT<br>ACCAGTACGGTGGCGGTAACTCTGCACTTGCTCTGC<br>AAACTGATGCCCGTAACTCTGACTTGACTATTACCCA<br>GCATGGCGGCGGTAATGGTGCAGATGTTGGTCAGG<br>GCTCAGATGACAGCTCAATCGATCTGACCCAACGTG<br>GCTTCGGTAACAGCGCTACTCTTGATCAGTGGAACG<br>GCAAAAATTCTGAAATGACGGTTAAACAGTTCGGTGG<br>TGGCAACGGTGCTGCAGTTGACCAGACTGCATCTAA<br>CTCCTCCGTCAACGTGACTCAGGTTGGCTTTGGTAA<br>CAACGCGACCGCTCATCAGTACTAA |

**Supplementary Figure 3. Extended Hel-CsgA hydrogel SEM imaging.**

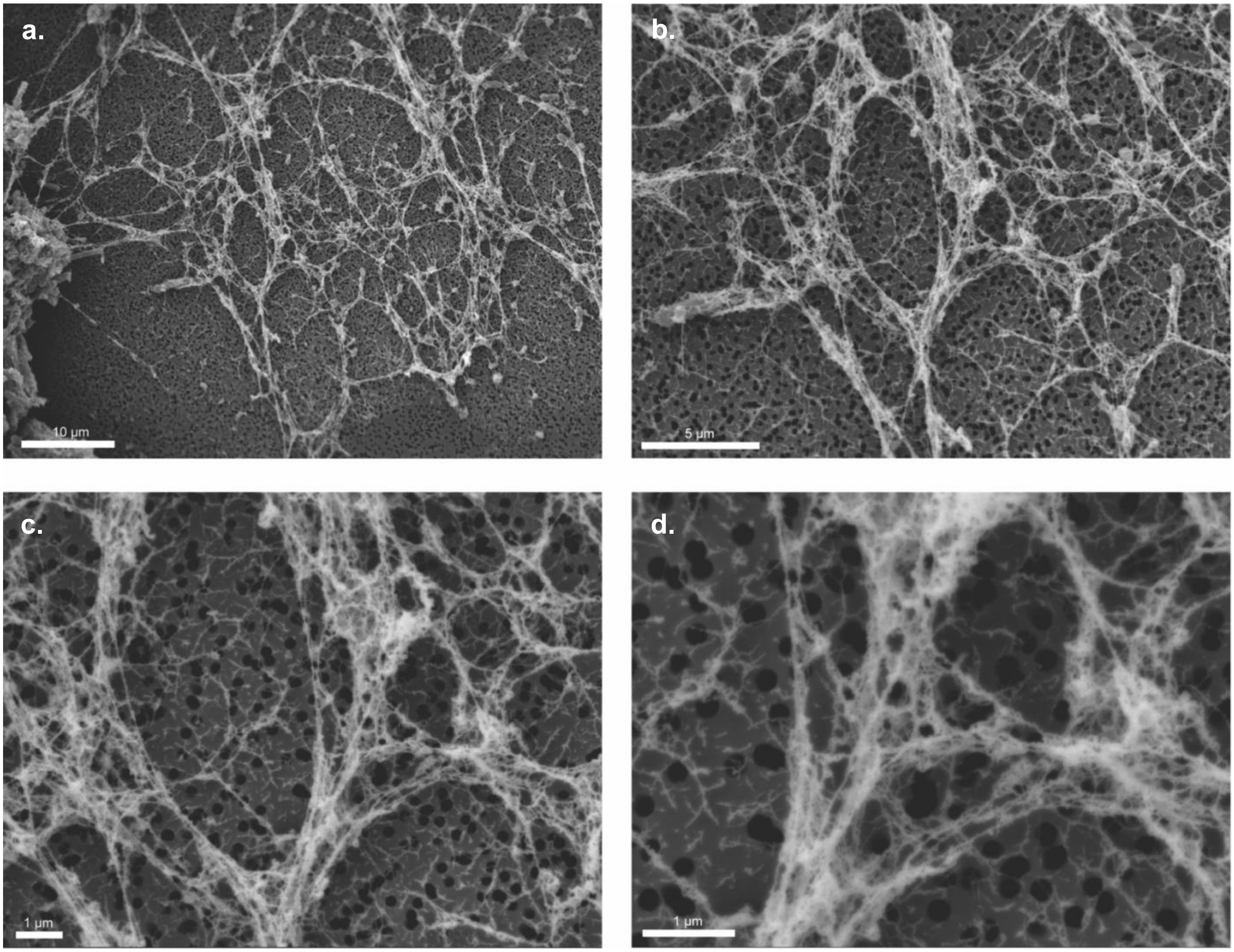

**Supplementary Figure 4. Extended CFU chart with controls.**

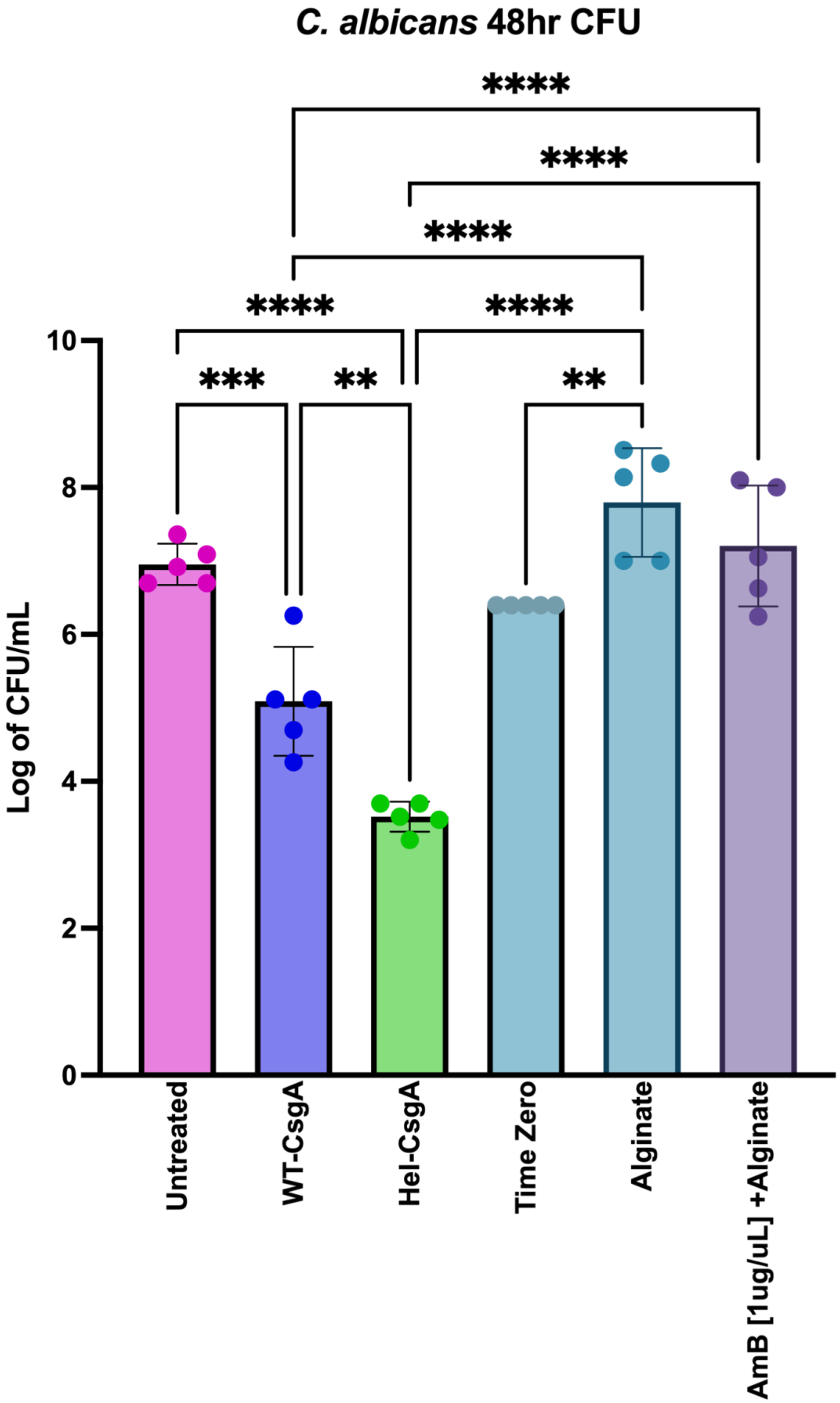

**Supplementary Figure 5. SEM Alginate fungal challenge imaging.**

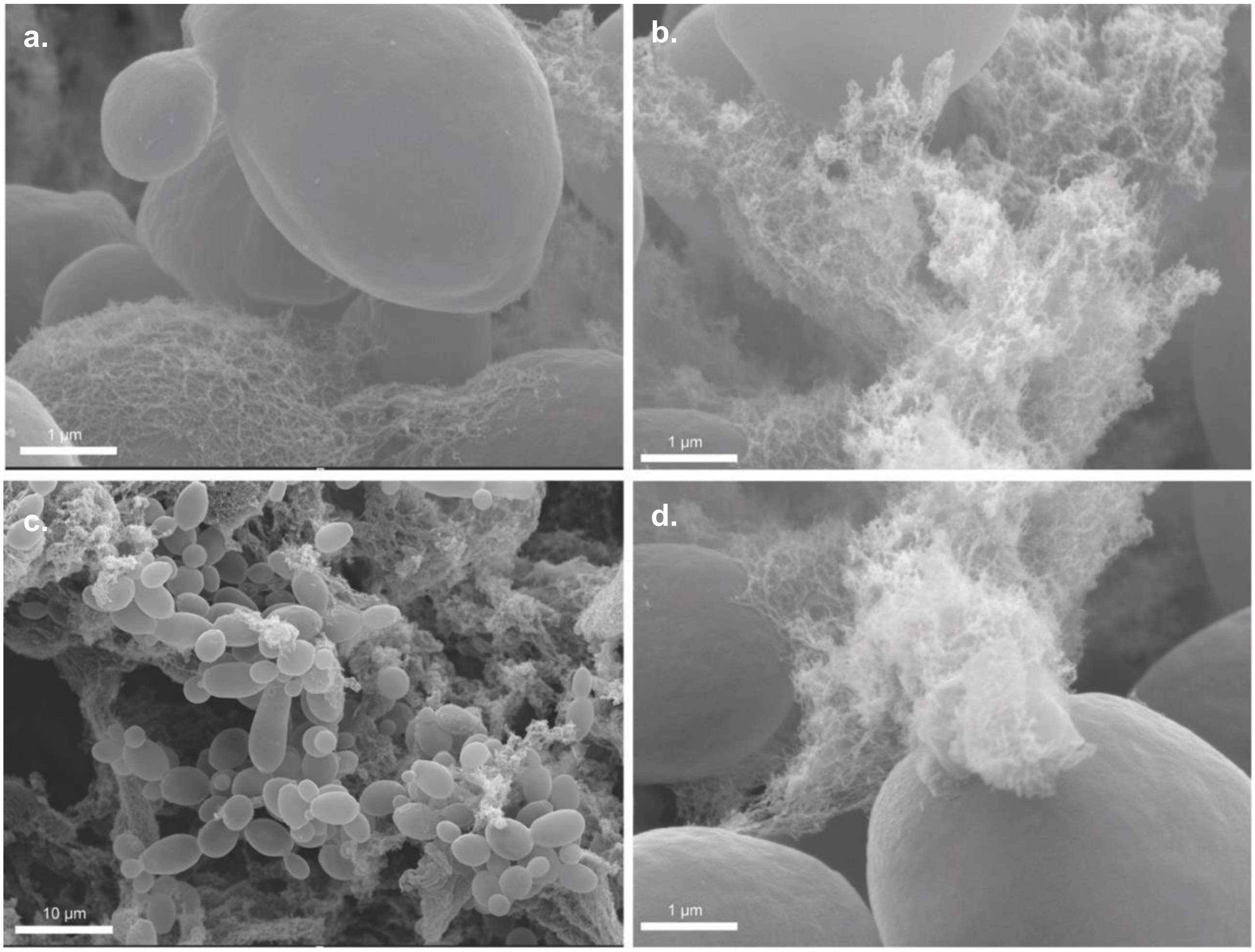

**Supplementary Figure 6. Extended SEM wtCsgA fungal challenge imaging.**

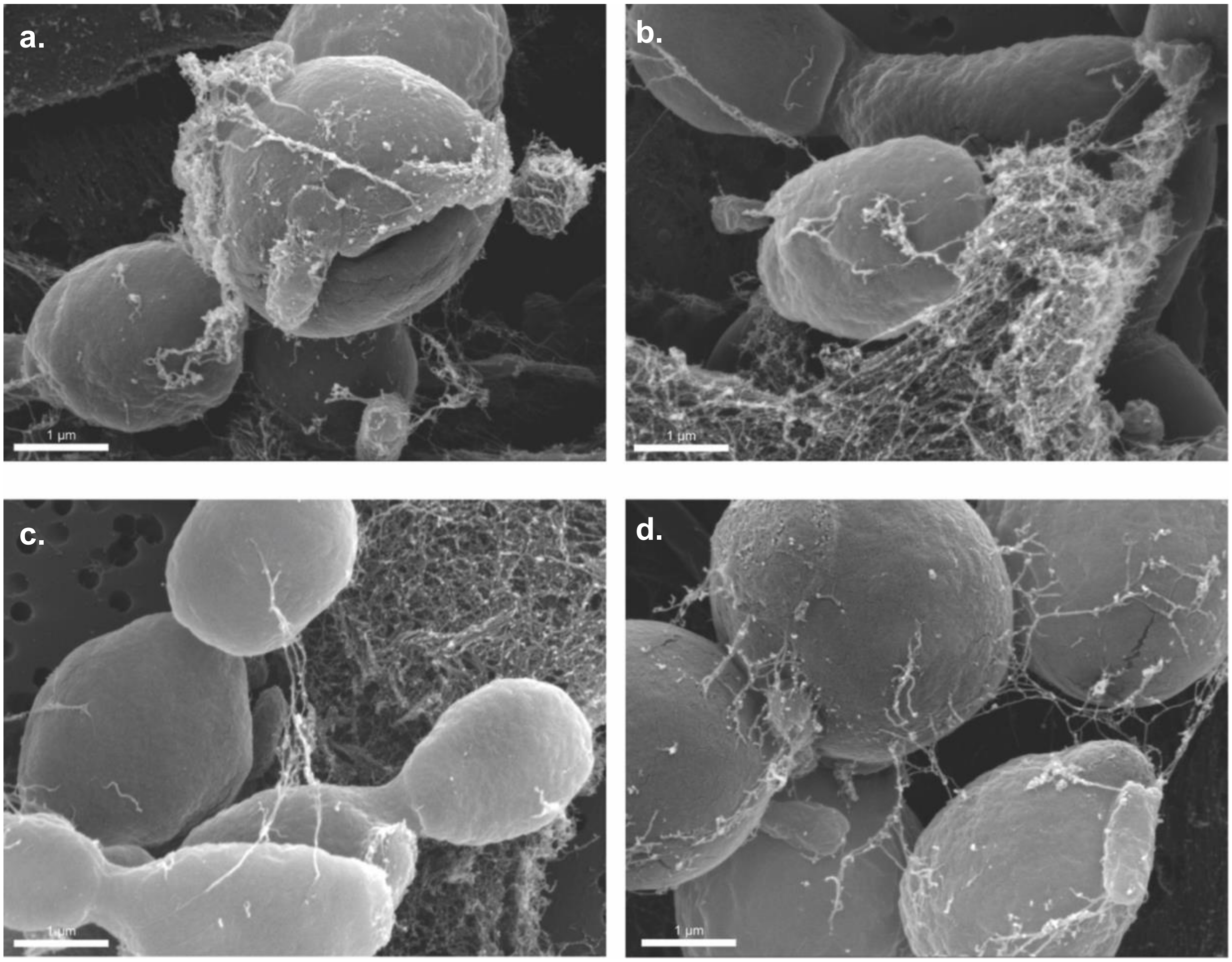

**Supplementary Figure 7. Extended SEM Hel-CsgA fungal challenge imaging.**

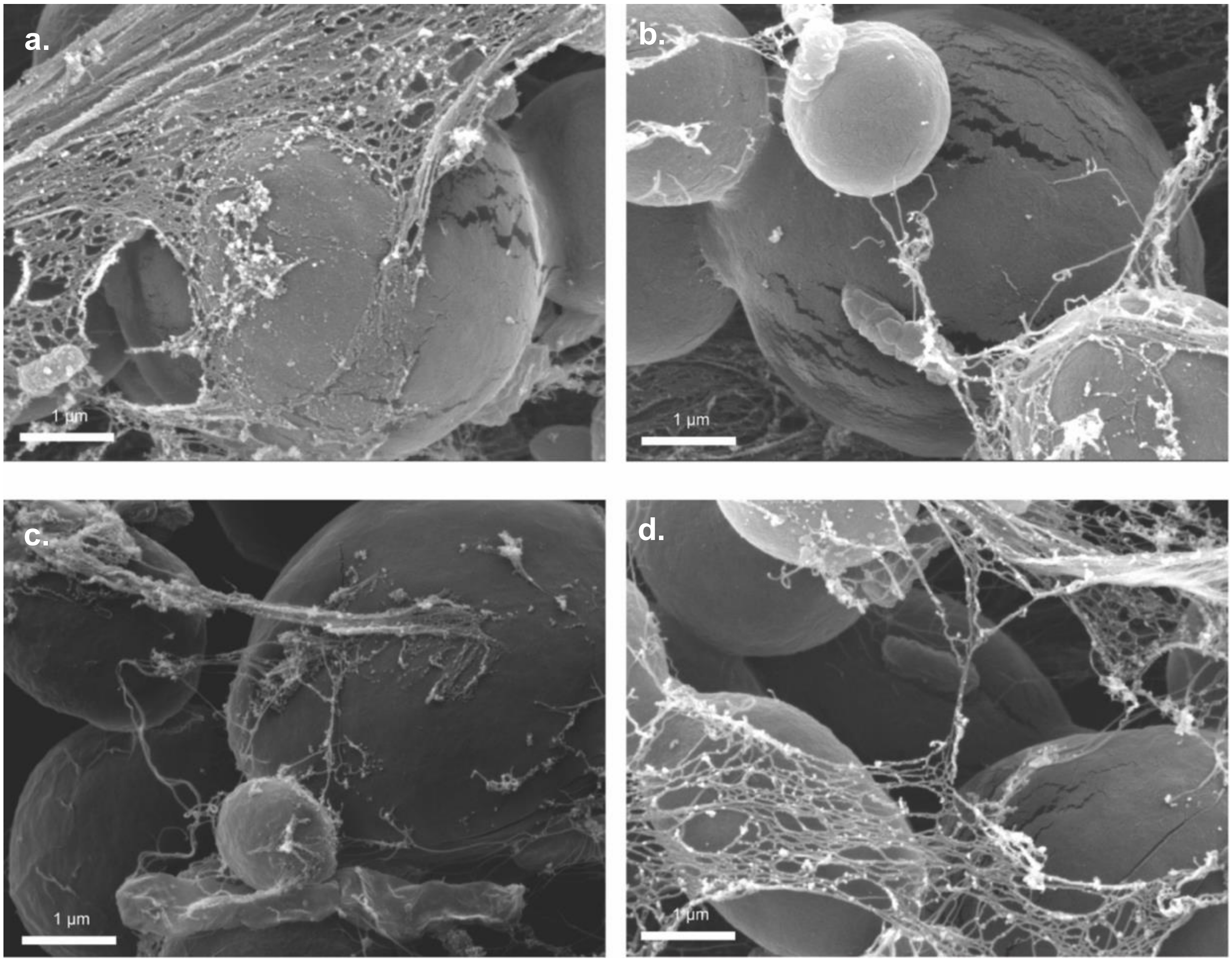

**Supplementary Figure 8. 3D Biomaterial Print Replicates (Rows: (a)untreated (b)wtCsgA, and (c)Hel-CsgA)**

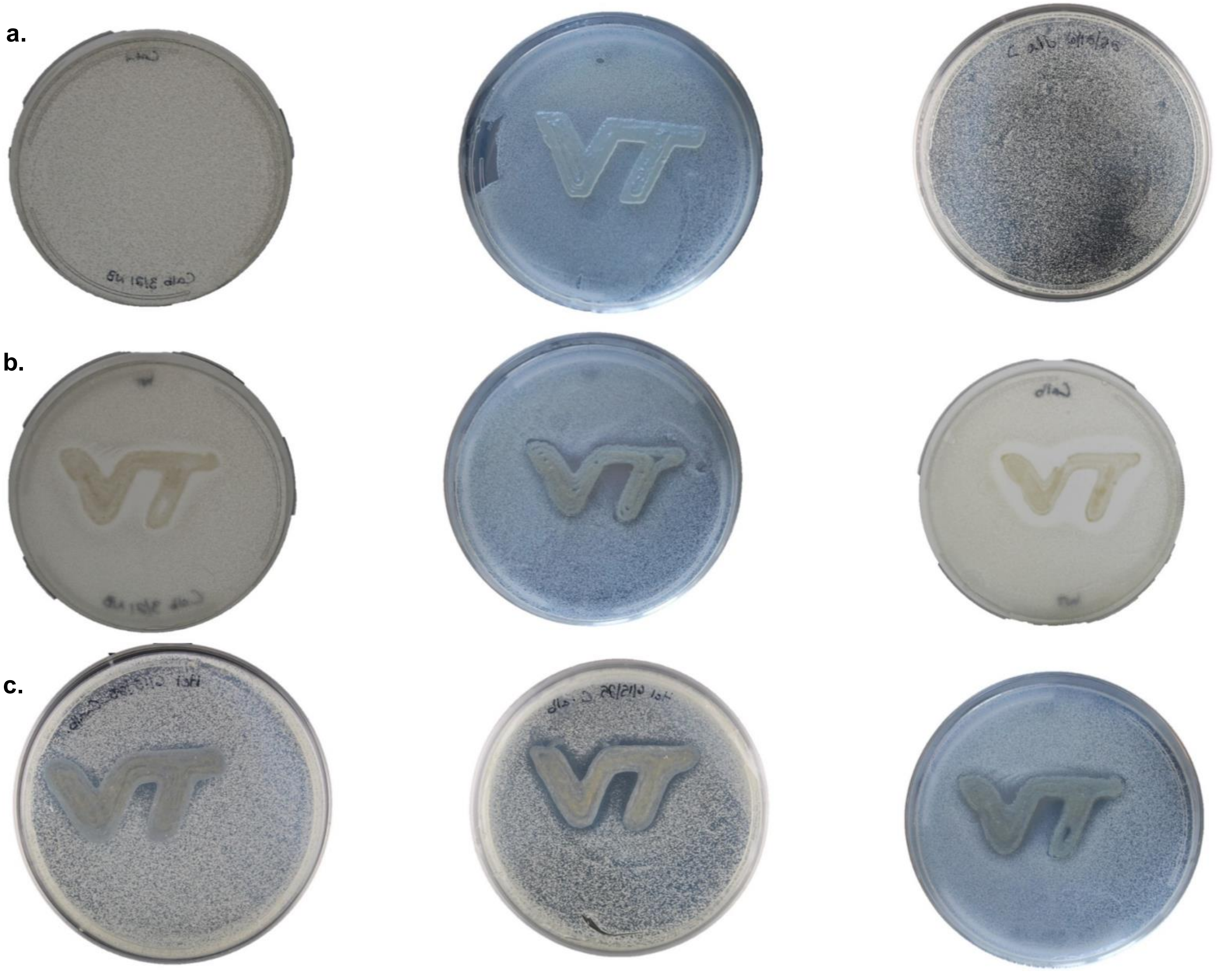

**Supplementary Figure 9. Extended Propidium Iodide assay.**

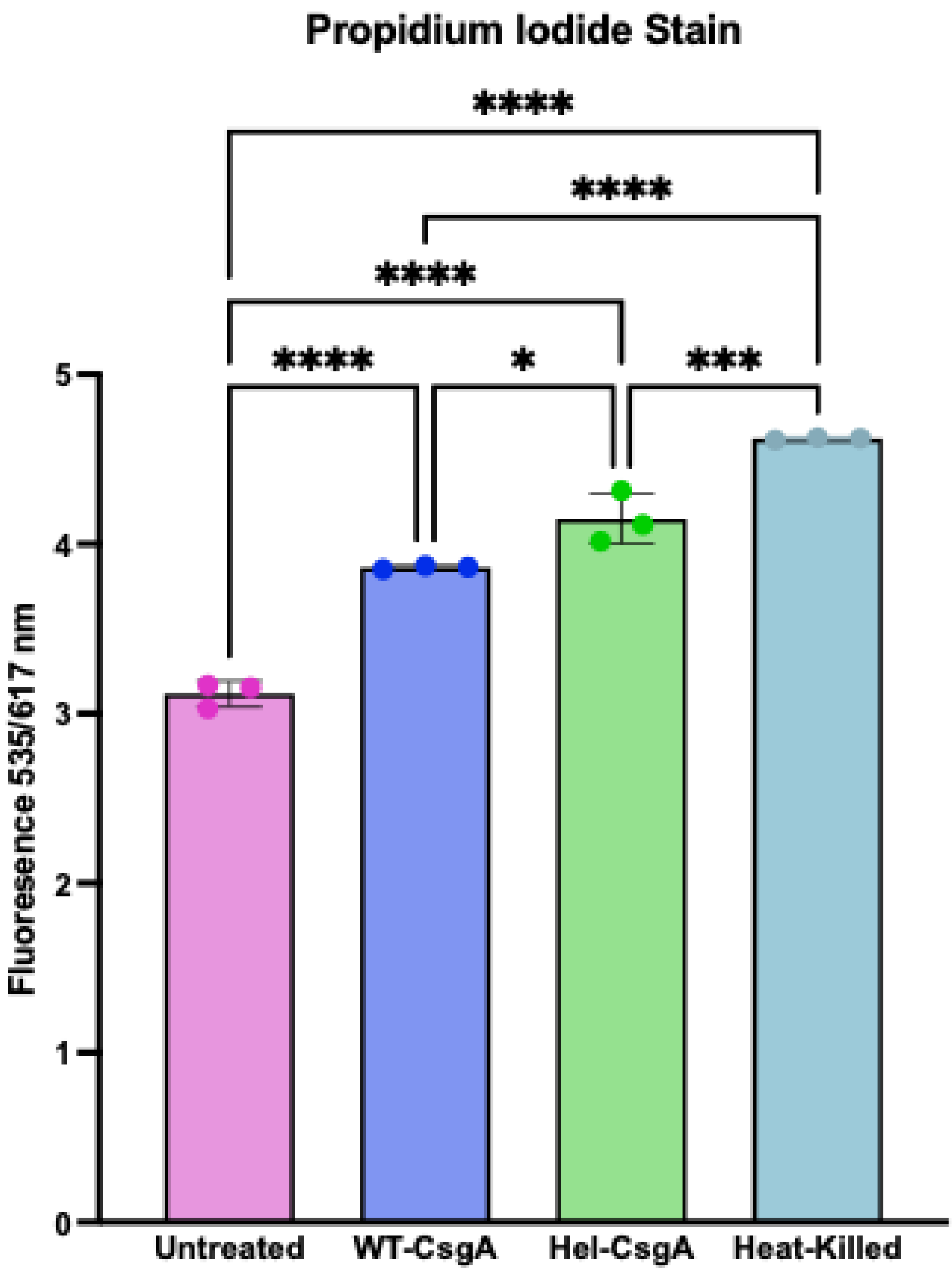

**Supplementary Figure 10. Extended Nile Red assay.**

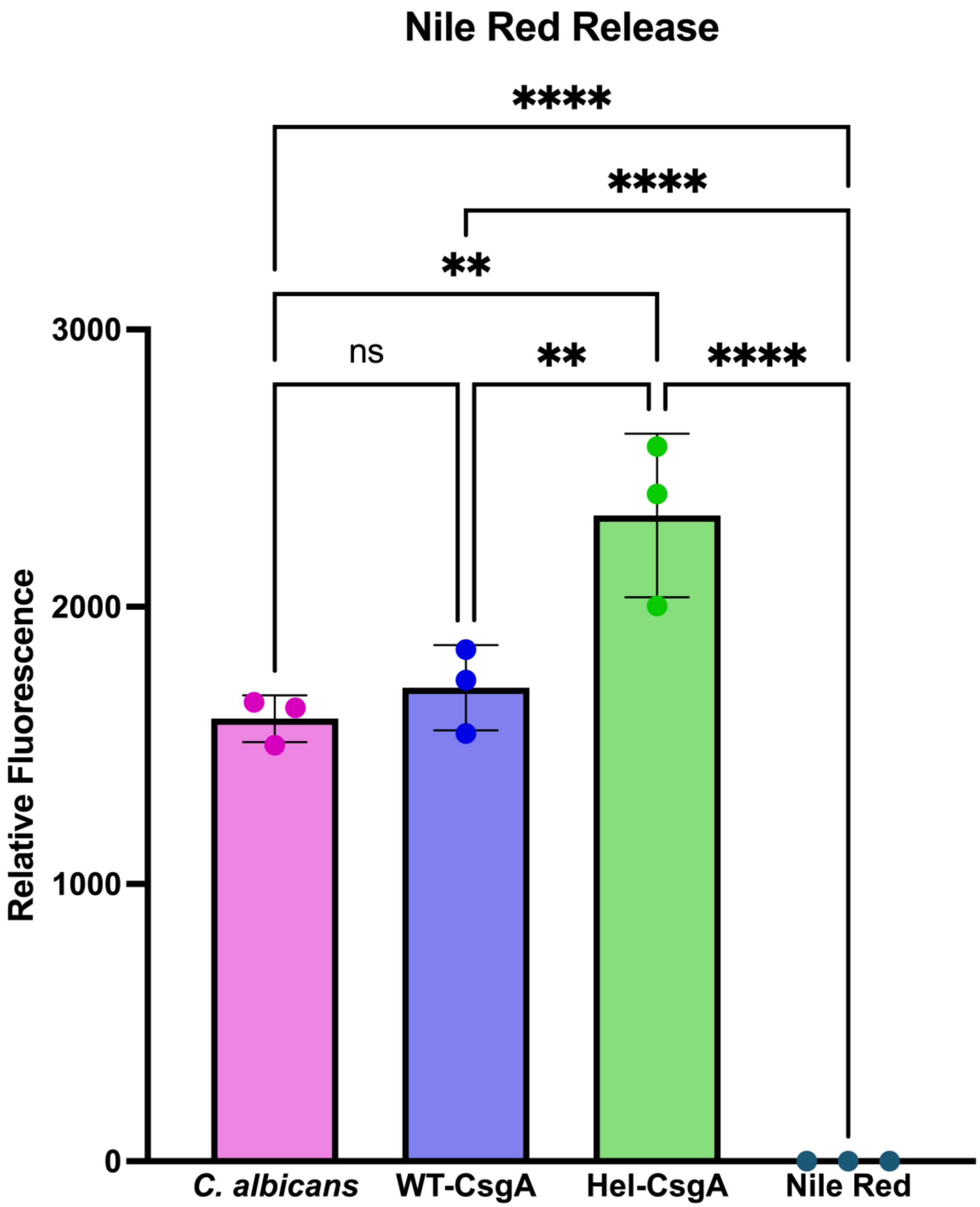

**Supplementary Table 2. Physiochemical properties of Hel- and WT-CsgA.**

| *Protein* | Molecular Weight (kDa) | Isoelectric Point (Pi) | Net Charge at pH 7.4 | Gravity |
|---|---|---|---|---|
| *WT-CsgA* | 13.1 | 4.528 | -6.32 | -0.718 |
| *Hel-CsgA* | 18.7 | 5.145 | -5.88 | -0.648 |

**Supplementary Table 3: wtCsgA amino acid composition.**

| Name | 3-Letter | Class | Number | Count (%) |
|---|---|---|---|---|
| **Alanine** | Ala | Hydrophobic | 10 | 7.63% |
| **Arginine** | Arg | Basic | 2 | 1.53% |
| **Asparagine** | Asn | Hydrophilic | 16 | 12.21% |
| **Aspartic Acid** | Asp | Acidic | 8 | 6.11% |
| **Cysteine** | Cys | Hydrophilic | 0 | 0% |
| **Glutamic Acid** | Glu | Acidic | 2 | 1.53% |
| **Glutamine** | Gln | Hydrophilic | 11 | 8.40% |
| **Glycine** | Gly | Hydrophilic | 28 | 21.37% |
| **Histidine** | His | Basic | 3 | 2.29% |
| **Isoleucine** | Ile | Hydrophobic | 3 | 2.29% |
| **Leucine** | Leu | Hydrophobic | 6 | 4.58% |
| **Lysine** | Lys | Basic | 2 | 1.53% |
| **Methionine** | Met | Hydrophobic | 1 | 0.76% |
| **Phenylalanine** | Phe | Hydrophobic | 3 | 2.29% |
| **Proline** | Pro | Hydrophobic | 2 | 1.53% |
| **Serine** | Ser | Hydrophilic | 12 | 9.16% |
| **Threonine** | Thr | Hydrophilic | 9 | 6.87% |
| **Tryptophan** | Trp | Hydrophobic | 1 | 0.76% |
| **Tyrosine** | Tyr | Hydrophobic | 4 | 3.05% |
| **Valine** | Val | Hydrophobic | 8 | 6.11% |

**Supplementary Table 4: Hel-CsgA amino acid composition.**

| Name | 3-Letter | Class | Number | Count (%) |
|---|---|---|---|---|
| **Alanine** | Ala | Hydrophobic | 12 | 6.42% |
| **Arginine** | Arg | Basic | 4 | 2.14% |
| **Asparagine** | Asn | Hydrophilic | 20 | 10.7% |
| **Aspartic Acid** | Asp | Acidic | 10 | 5.35% |
| **Cysteine** | Cys | Hydrophilic | 6 | 3.21% |
| **Glutamic Acid** | Glu | Acidic | 4 | 2.14% |
| **Glutamine** | Gln | Hydrophilic | 11 | 5.88% |
| **Glycine** | Gly | Hydrophilic | 43 | 22.99% |
| **Histidine** | His | Basic | 4 | 2.14% |
| **Isoleucine** | Ile | Hydrophobic | 4 | 2.14% |
| **Leucine** | Leu | Hydrophobic | 7 | 3.74% |
| **Lysine** | Lys | Basic | 5 | 2.67% |
| **Methionine** | Met | Hydrophobic | 1 | 0.53% |
| **Phenylalanine** | Phe | Hydrophobic | 4 | 2.14% |
| **Proline** | Pro | Hydrophobic | 2 | 1.07% |
| **Serine** | Ser | Hydrophilic | 19 | 10.16% |
| **Threonine** | Thr | Hydrophilic | 11 | 5.88% |
| **Tryptophan** | Trp | Hydrophobic | 3 | 1.6% |
| **Tyrosine** | Tyr | Hydrophobic | 6 | 3.21% |
| **Valine** | Val | Hydrophobic | 11 | 5.88% |

**Supplementary Figure 11. Extended *Ex vivo C. albicans* no-treatment imaging.**

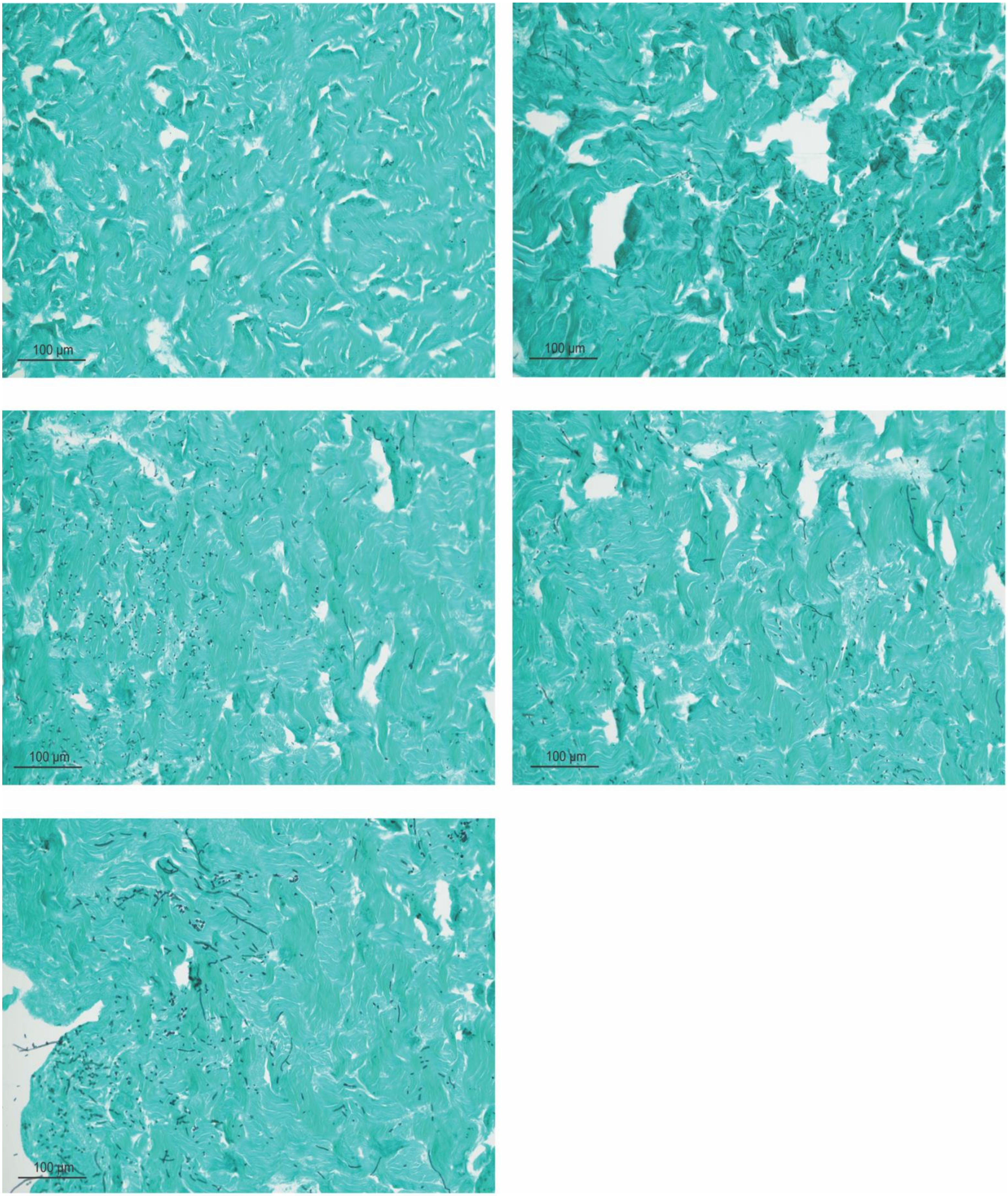

**Supplementary Figure 12. Extended *Ex vivo C. albicans* wtCsgA treatment Imaging.**

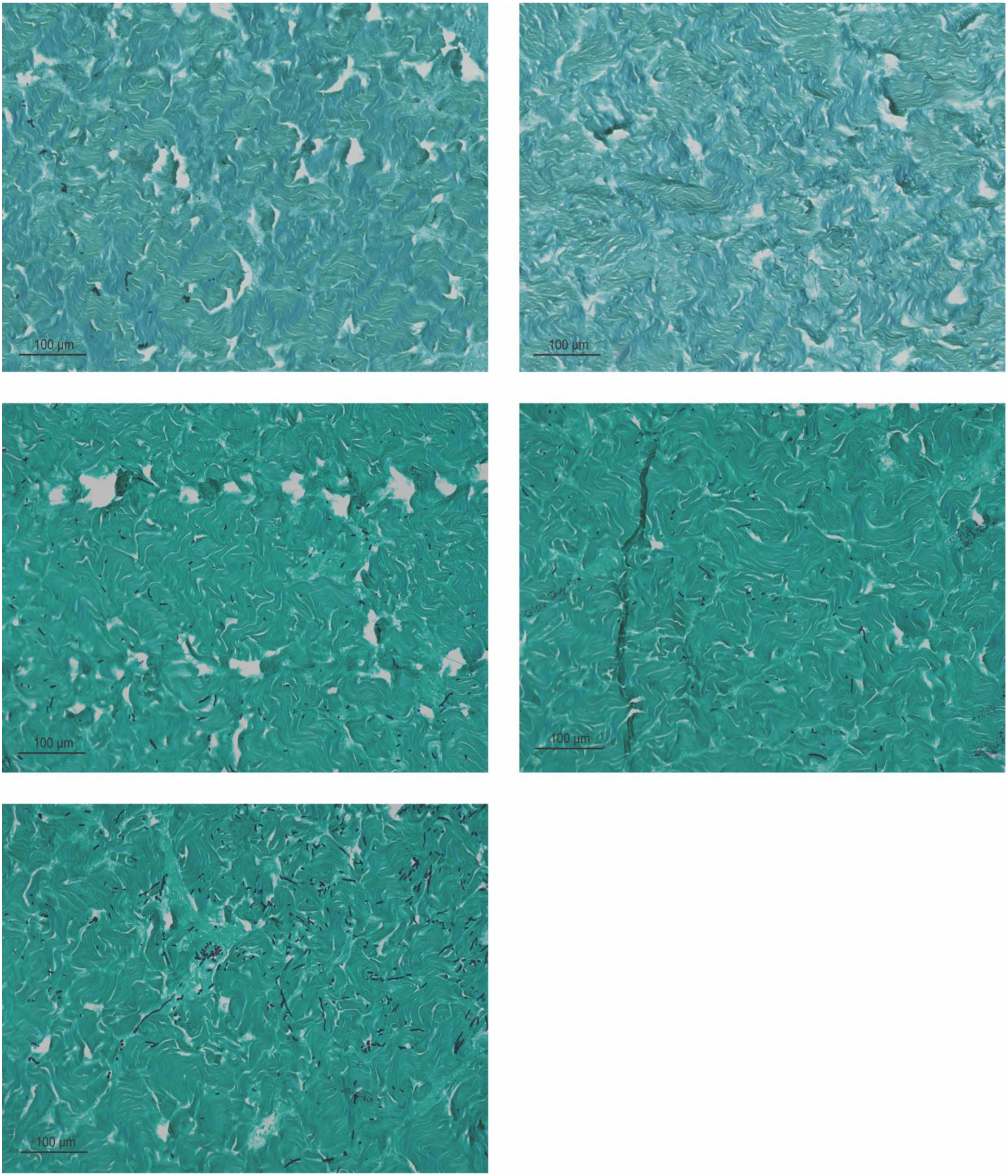

**Supplementary Figure 13. Extended *Ex vivo C. albicans* Hel-CsgA treatment Imaging.**

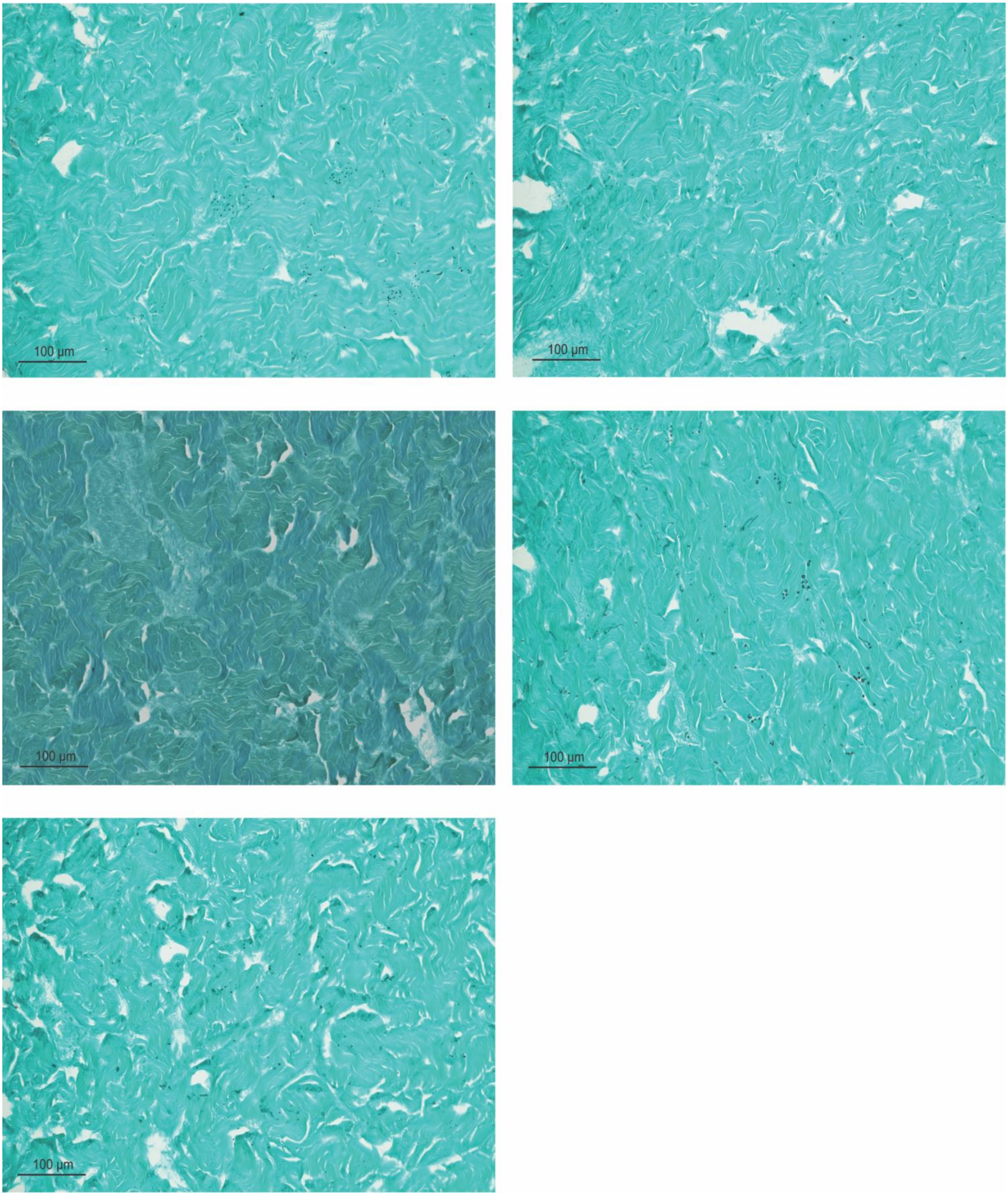
